# Human single-neuron activity is modulated by intracranial theta burst stimulation of the basolateral amygdala

**DOI:** 10.1101/2024.11.11.622161

**Authors:** Justin M. Campbell, Rhiannon L. Cowan, Krista L. Wahlstrom, Martina K. Hollearn, Dylan Jensen, Tyler Davis, Shervin Rahimpour, Ben Shofty, Amir Arain, John D. Rolston, Stephan Hamann, Shuo Wang, Lawrence N. Eisenman, James Swift, Tao Xie, Peter Brunner, Joseph R. Manns, Cory S. Inman, Elliot H. Smith, Jon T. Willie

**Affiliations:** Interdepartmental Program in Neuroscience, University of Utah, Salt Lake City, UT, USA; Department of Neurosurgery, University of Utah, Salt Lake City, UT, USA; Department of Psychology, University of Utah, Salt Lake City, UT, USA; Department of Neurology, University of Utah, Salt Lake City, UT, USA; Department of Neurosurgery, Brigham and Women’s Hospital, Boston, MA, USA; Department of Psychology, Emory University, Atlanta, GA, USA; Department of Radiology, Washington University School of Medicine, St. Louis, MO, USA; Department of Neurology, Washington University School of Medicine, St. Louis, MO, USA; Department of Neurological Surgery, Washington University School of Medicine, St. Louis, MO, USA; National Center for Adaptive Neurotechnologies, St. Louis, MO, USA; Senior author

**Keywords:** Intracranial EEG, Single-unit, Theta burst stimulation, Modulation, Amygdala

## Abstract

Direct electrical stimulation of the human brain has been used for numerous clinical and scientific applications. At present, however, little is known about how intracranial stimulation affects activity at the microscale. In this study, we recorded intracranial EEG data from a cohort of patients with medically refractory epilepsy as they completed a visual recognition memory task. During the memory task, brief trains of intracranial theta burst stimulation (TBS) were delivered to the basolateral amygdala (BLA). Using simultaneous microelectrode recordings, we isolated neurons in the hippocampus, amygdala, orbitofrontal cortex, and anterior cingulate cortex and tested whether stimulation enhanced or suppressed firing rates. Additionally, we characterized the properties of modulated neurons, clustered presumed excitatory and inhibitory neurons by waveform morphology, and examined the extent to which modulation affected memory task performance. We observed a subset of neurons (∼30%) whose firing rate was modulated by TBS, exhibiting highly heterogeneous responses with respect to onset latency, duration, and direction of effect. Notably, location and baseline activity predicted which neurons were most susceptible to modulation, although the impact of this neuronal modulation on memory remains unclear. These findings advance our limited understanding of how focal electrical fields influence neuronal firing at the single-cell level.

**Highlights:**

Individual neurons in the human brain were responsive to theta burst stimulation
Basolateral amygdala stimulation preferentially modulated neurons in the hippocampus, orbitofrontal cortex, and amygdala
Excitatory and inhibitory neurons were modulated equally
Neurons modulated by stimulation tended to have greater baseline firing rates
Duration, onset, and valence of neuronal modulation were heterogeneous

## Introduction

The amygdala is a highly connected cluster of nuclei with input from multiple sensory modalities and vast projections to distributed cortical and subcortical regions involved in autonomic regulation and cognition (Kim et al., 2011; McDonald and Mott, 2017; Roy et al., 2009; Stein et al., 2007). Numerous studies have described the amygdala’s capacity to facilitate the encoding of long-lasting emotional memories (Dolcos et al., 2004; Hermans et al., 2014; LaBar and Cabeza, 2006; Phelps, 2006, 2004; Phelps and LeDoux, 2005; Qasim et al., 2023; Richardson et al., 2004; Roozendaal et al., 2009; Schwabe et al., 2013; Zheng et al., 2017). Specific oscillatory dynamics in the amygdalohippocampal circuit are increasingly understood to be essential in prioritizing the encoding of these salient memories (Qasim et al., 2023; Zheng et al., 2019, 2017).

Recently, direct electrical stimulation of the basolateral complex of the amygdala (BLA) in humans has revealed a more generalized ability to enhance declarative memory irrespective of the emotional valence (Inman et al., 2018), likely by promoting synaptic plasticity-related processes underlying memory consolidation in the hippocampus and medial temporal lobe (McGaugh, 2013, 2004; McGaugh et al., 2002; Roesler et al., 2021). These pro-memory effects were achieved with rhythmic theta-burst stimulation (TBS), which is known to induce long-term potentiation (LTP), a key mechanism in memory formation (Larson and Munkácsy, 2015). Emerging evidence suggests that intracranial TBS may also enhance memory specificity (Titiz et al., 2017), evoke theta-frequency oscillations (Solomon et al., 2021), and facilitate short-term plasticity in local field potential recordings (Herrero et al., 2021; Huang et al., 2024). However, the extent to which exogenous TBS modulates activity at the single-cell level and whether this modulation is associated with memory performance remains poorly understood.

Here, we address this knowledge gap by conducting simultaneous microelectrode recordings from prefrontal and medial temporal structures during a memory task in which intracranial TBS was applied to the BLA. To characterize neuronal modulation, we contrasted the trial-averaged, peri-stimulation firing rates across stim and no-stim conditions; neurons were further classified based on their location, baseline activity, and direction of effect (enhancement vs. suppression). We predicted that intracranial TBS of the BLA would modulate spiking activity within highly connected regions (e.g., hippocampus) and improve memory task performance.

## Methods

### Participants

We report results from a cohort of 23 patients with medically refractory epilepsy who underwent stereoelectroencephalography to localize epileptogenic foci (74% female, 19– 66 years old). All patients were age 18+ and able to provide informed consent. No exclusion was made concerning a patient’s sex, gender, race, ethnicity, or socioeconomic status. Surgeries were performed at the University of Utah in Salt Lake City, UT (n = 10) and Barnes-Jewish Hospital in St. Louis, MO (n = 13). Patients were monitored continuously by a clinical team during their post-operative hospital course. Each patient signed a written informed consent form before participation in the research study; all study procedures were approved by the Institutional Review Board at the University of Utah (IRB 00144045, IRB 00114691) and Washington University (IRB 202104033).

### Electrode Placement and Localization

Numbers and trajectories of stereoelectroencephalography electrode placements were determined case-by-case and solely derived from clinical considerations during a multidisciplinary case conference without reference to this research program. Each patient was implanted with clinical macroelectrodes and 1–3 Behnke-Fried depth electrodes (Ad-Tech Medical Instrument Corporation, Oak Creek, WI), which contained both macro- and microelectrode contacts (eight 40 µm diameter microwires and one unshielded reference wire) for recording local field potentials and extracellular action potentials, respectively (*Figure 1A*). To localize electrodes, we leveraged the open-access *Localize Electrodes Graphical User Interface (LeGUI)* (Davis et al., 2021) software developed by our group, which performs coregistration of pre-operative MRI and post-operative CT sequences, image processing, normalization to standard anatomical templates, and automated electrode detection.

**Figure 1:**
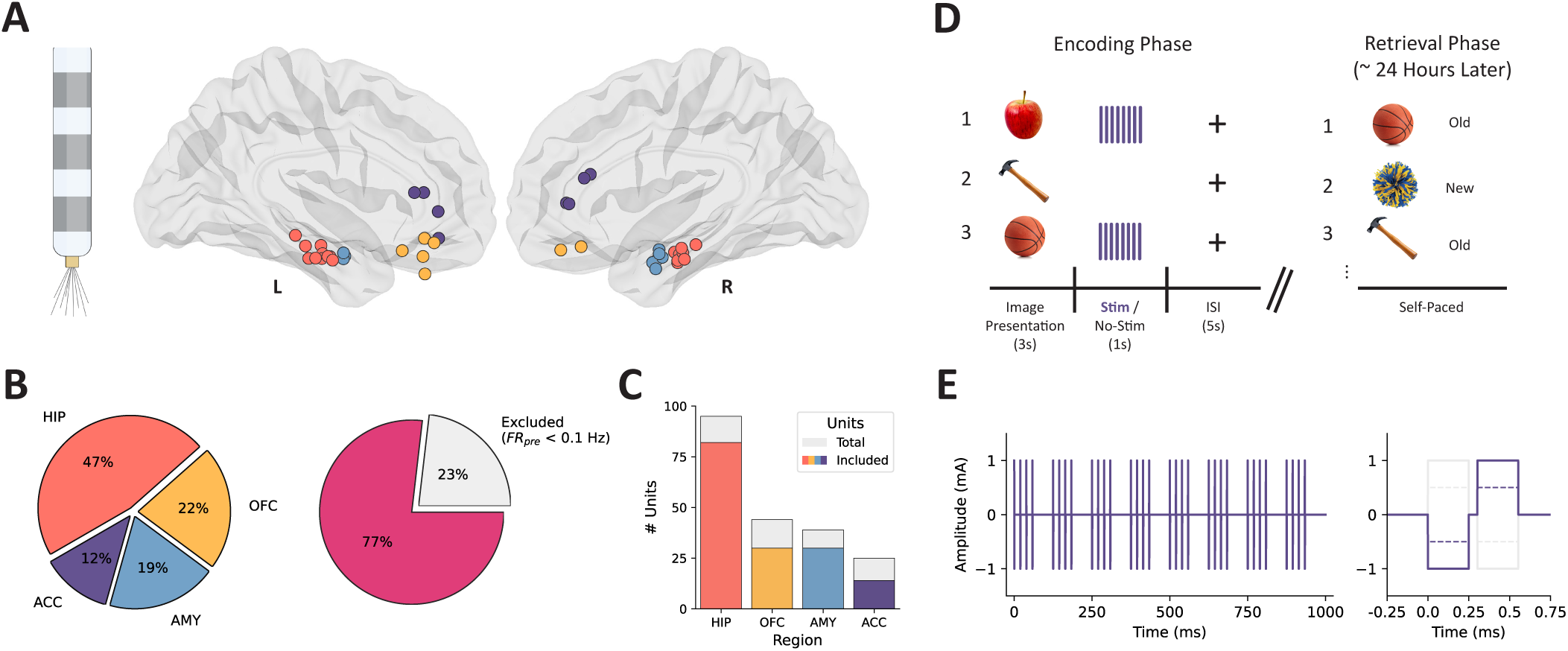
Microelectrode locations, unit counts, and experimental design. (A) Behnke Fried-style macro/micro depth electrode (left) and microelectrode bundle locations projected in MNI space (right). (B) The proportion of units recorded from each brain area (left) and the proportion of units that met the criteria for inclusion in analyses (average pre-trial baseline firing rate ≥ 0.1 Hz) (right). (C) Counts of total (grey) and included (colored) units within each region. (D) Intracranial recording and stimulation took place in the context of a two-phase (encoding, retrieval) visual recognition memory task. A series of neutral valence images were shown (3 s), half of which were followed by direct electrical stimulation (1 s). Retrieval memory was tested during a self-paced task ∼24 hours later. (E) Simulated theta-burst stimulation trace (left) and individual stimulation pulse (right); charge-balanced, bipolar, biphasic rectangular pulses were delivered over a 1 s period. HIP = hippocampus (coral), OFC = orbitofrontal cortex (yellow), AMY = amygdala (blue), ACC = anterior cingulate cortex (purple).

### Intracranial Electrophysiology

Neurophysiological data were recorded at both hospitals using a neural signal processor (Blackrock Microsystems, Salt Lake City, UT; Nihon Koden USA, Irvine, CA) sampling at 30 kHz. Microelectrode contacts were locally referenced to a low-impedance microwire near the recording wires. Macroelectrode contacts were referenced to an intracranial contact located within the white matter with minimal activity, per recommended practices (Mercier et al., 2022).

### Experimental Design

Patients completed a visual recognition memory task previously employed by our group to characterize the effects of basolateral amygdala stimulation upon memory consolidation (Inman et al., 2018). The memory task consisted of an encoding session, during which a series of neutral valence images were presented, and a self-paced retrieval session ∼24 hours post-encoding wherein patients were asked to indicate whether each image onscreen was old (previously shown) or new (unseen) (*Figure 2A*). Data were aggregated across four highly similar experimental paradigms with minor differences in the content of visual stimuli, number of trials, stimulation parameters, and testing intervals. Each encoding session consisted of 160 or 320 trials wherein an image was presented on screen for 3 s, followed by a ∼6 s interstimulus interval (fixation cross on screen).

**Figure 2:**
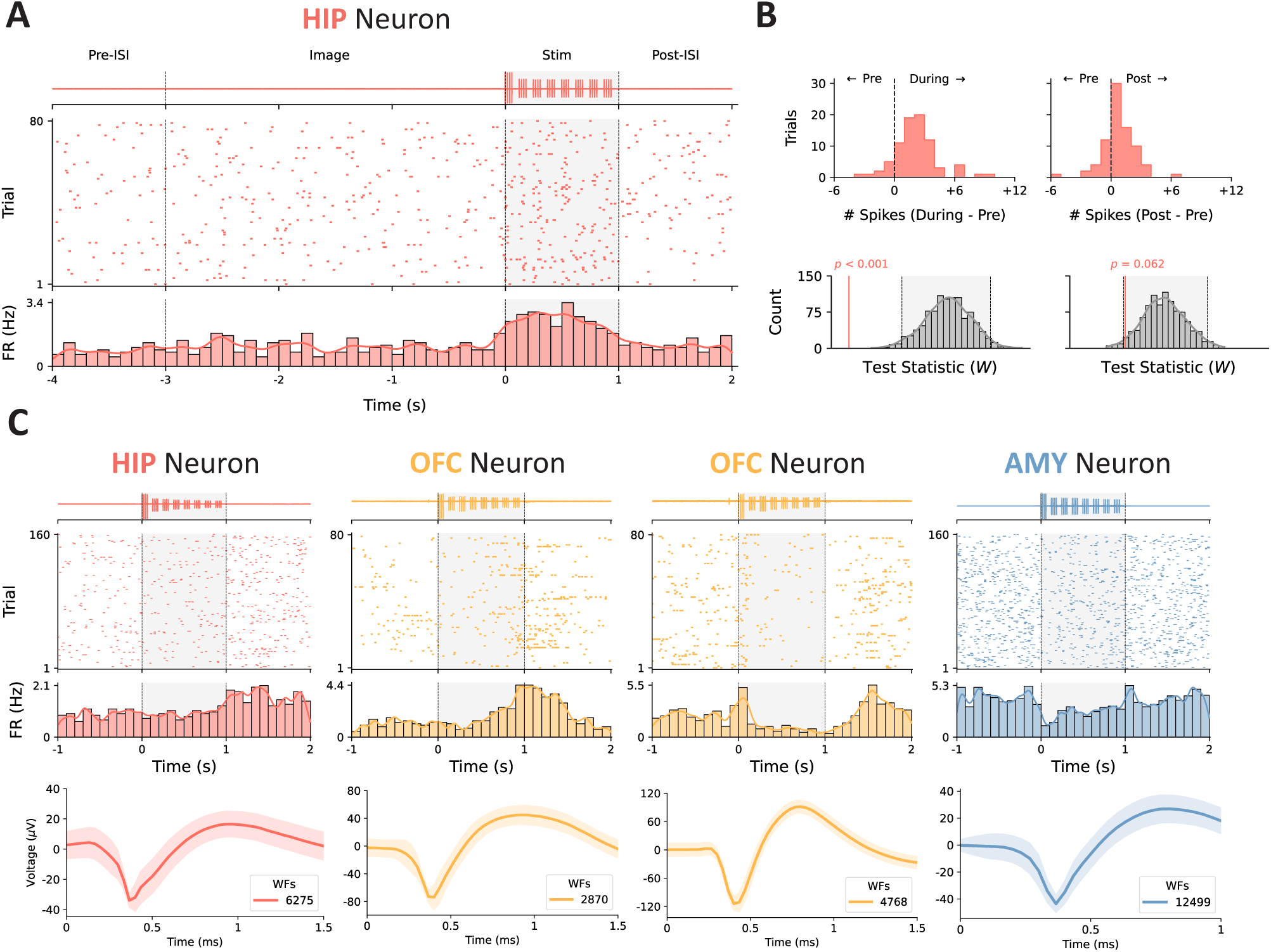
Example raster plots depicting heterogeneous responses to stimulation. (A) Representative example of modulation during stimulation. The highpass-filtered, trial-averaged LFP from the corresponding microwire is shown (top) above the spike raster for an example unit located in the hippocampus (middle); the grey shaded region depicts the duration of stimulation with onset at t = 0. The average firing rate across trials was estimated by convolving the binned spike counts (100 ms bins) with a Gaussian kernel (bottom). (B) The difference in the number of spikes in the 1 s peri-stimulation epochs for each trial is shown (top). We subsequently performed a Wilcoxon signed-rank test on the during- and post-stimulation spike counts for each trial vs. the pre-trial baseline and compared the empirical test statistic against a null distribution generated by shuffling the epoch labels 1,000 times (bottom); the grey-shaded region represents the distribution containing 95% of observed values. (C) Some units (left, left-middle) exhibited increased firing rates, whereas others (right-middle, right) had their firing suppressed. The temporal dynamics of the firing rate modulation (e.g., onset, duration) were highly variable across units. The averaged waveform for each of the visualized units is shown below its corresponding peri-stimulation raster plot (WFs = waveforms); the shaded region represents standard deviation across waveforms.

### Spike Detection and Sorting

Microelectrode data were first filtered between 250–500 Hz with a zero-phase lag bandpass filter and re-thresholded offline at -3.5 times the root mean square of the signal to identify spike waveforms. Units were isolated during a semi-automated process within *Offline Sorter* (Plexon Inc, Dallas, TX) by applying the T-distribution expectation maximization method on the first three principal components of the detected waveforms (initial parameters: degrees of freedom multiplier = 4, initial number of units = 3) (Shoham et al., 2003). Finally, the waveform shapes, interspike interval distribution, consistency of firing, and isolation from other waveform clusters were manually inspected for further curation and removal of spurious, non-physiological threshold crossings that could represent stimulation artifacts.

### Single Unit Quality Metrics

We calculated several distinct metrics to characterize detected units’ properties and assess the quality of our spike sorting (*Figure S2*): the number of units detected per microelectrode bundle, the mean firing rate (Hz) for each unit, the percentage of interspike intervals < 3 ms, the coefficient of variation across each unit’s spike train, the average presence ratio of firing in 1s bins (proportion of bins which contain ≥ 1 spike), the ratio between the peak amplitude of the averaged waveform and its standard deviation, and the mean signal-to-noise ratio of the averaged waveform.

### Discrimination of Excitatory vs. Inhibitory Neurons

We calculated two metrics from the averaged waveform from each detected unit: the valley-to-peak-width (VP) and the peak half-width (PHW) (*Figure 4A*); previously, these two properties of waveform morphology have been used to discriminate pyramidal cells (excitatory) from interneurons (inhibitory) in human intracranial recordings (Peyrache et al., 2012). Next, we performed k-means clustering (n = 2 clusters) on the waveform metrics, in line with previous approaches to cell type classification.

**Figure 3:**
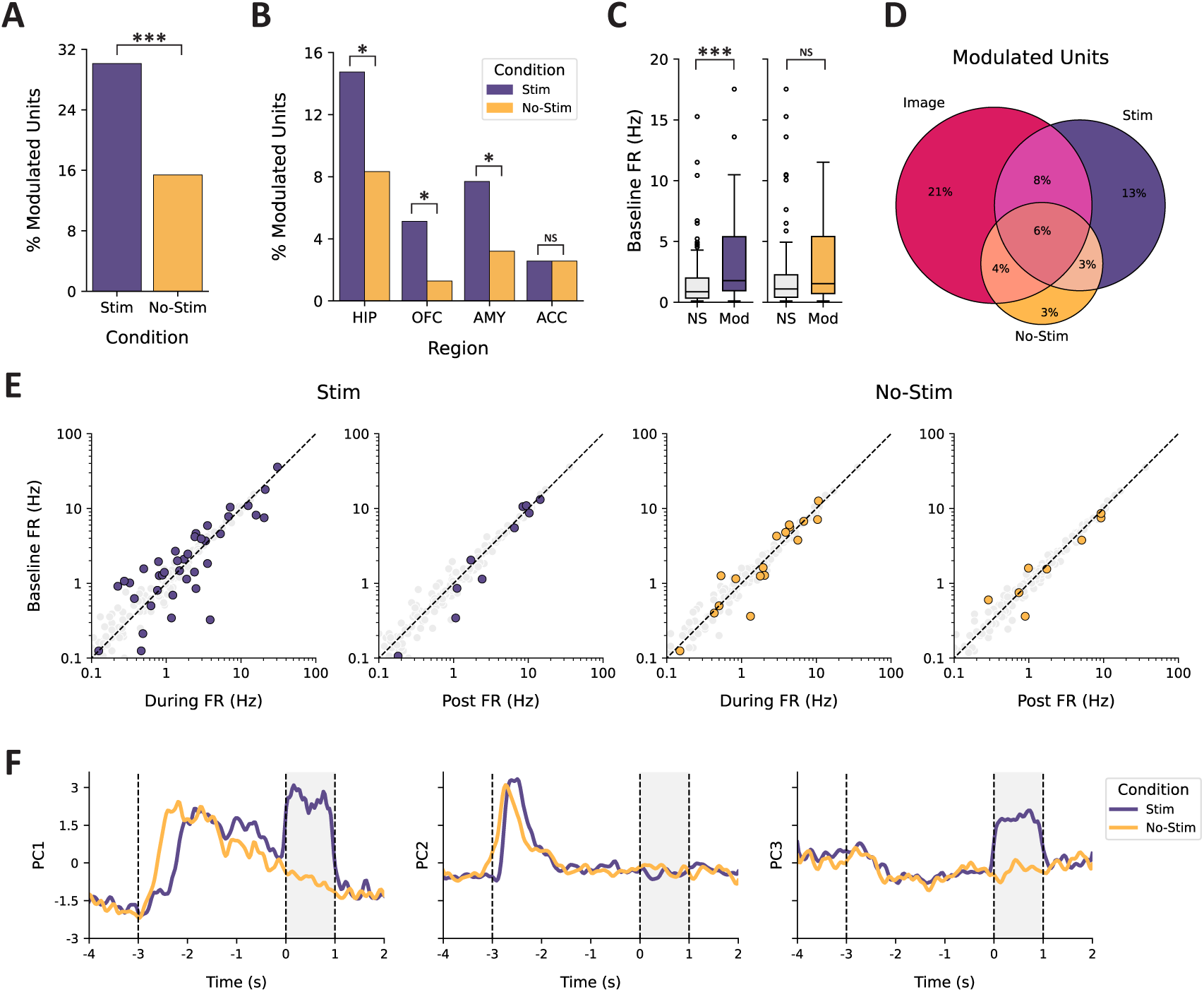
Characterization of modulation in neuronal firing rate. (A) Percent of modulated units observed across trials separated by stim (purple) vs. no-stim conditions (orange). (B) Percent of modulated units as a function of recording region. (C) Comparison of baseline firing rate in units separated by condition (stim vs. no-stim) and outcome (NS = not significant, Mod = modulated). (D) Venn diagram depicting the shared and independent proportions of units modulated by image onset (Image) and the two experimental conditions (stim vs. no-stim). (E) Scatterplot of pre-stimulation firing rate relative to the firing rate during the two contrast windows (during, post) for the stim (left) and no-stim (right) conditions. Modulated units are highlighted in purple (stim) or orange (no-stim), whereas units without a significant change are shown in grey. (F) Temporal dynamics of pseudo-population coactivity within each condition, represented by the first three principal components of the trial-averaged firing rates. The grey-shaded region depicts the duration of stimulation with onset at t = 0. Images were presented on screen for 3 s, with onset at t = -3. * *p* < 0.05, *** *p* < 0.001, NS = not significant.

**Figure 4:**
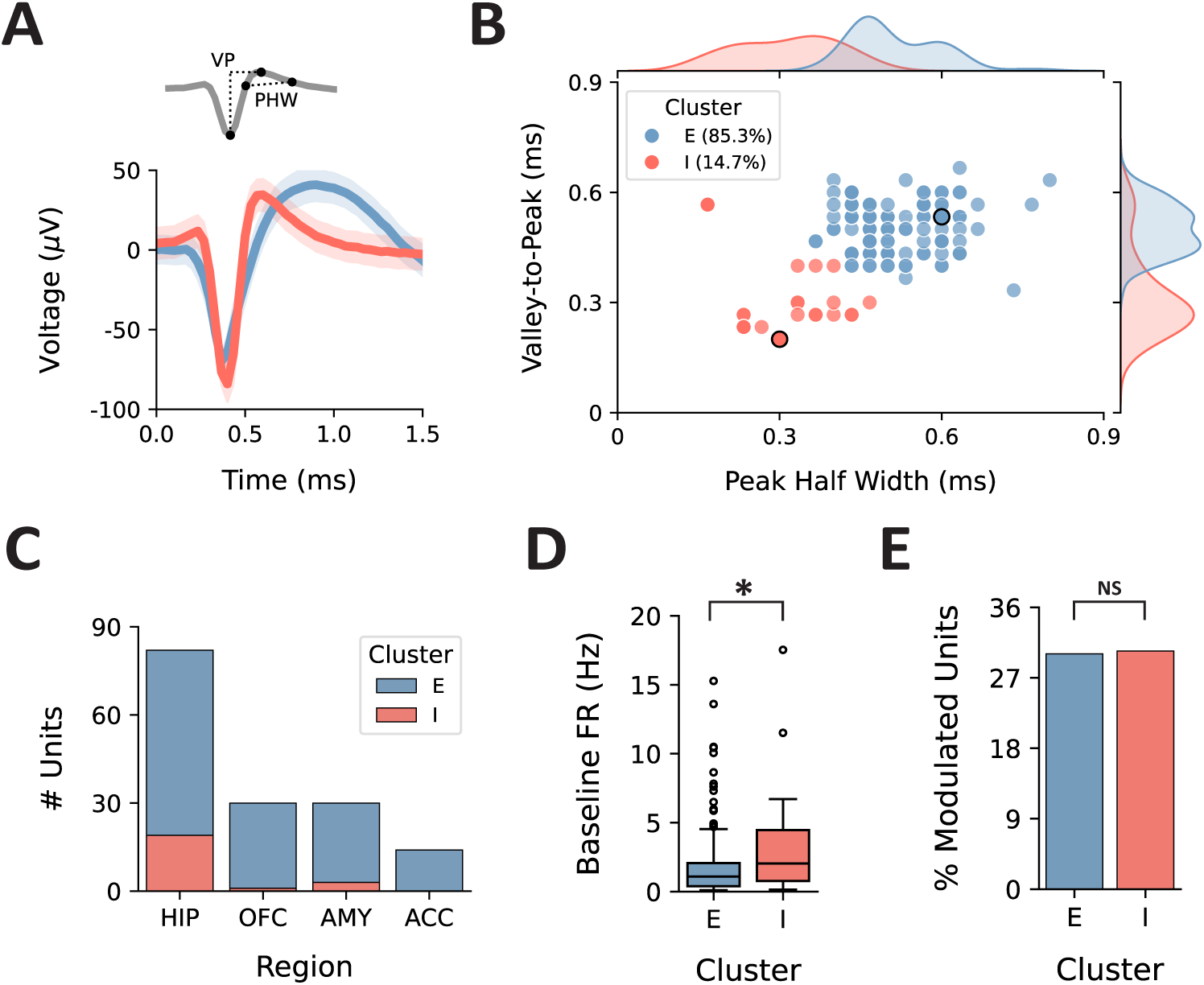
Separation of presumed excitatory and inhibitory neurons by waveform morphology. (A) Two metrics were calculated using the averaged waveforms for each detected unit: the valley-to-peak width (VP) and peak half-width (PHW). (B) Scatterplot of the relationship between VP and PHW; note that units with identical metrics are overlaid. Using k-means clustering, we identified two distinct response clusters, representing presumed excitatory (E, blue) and inhibitory (I, red) neurons. The units from which the example waveforms were taken are outlined in black. Probability distributions for each metric are shown along the axes. (C) Total number of units within each cluster, separated by region. (D) Comparison of baseline firing rates, separated by cluster. (E) Percent of modulated units in each cluster. * *p* < 0.05, NS = not significant.

### Intracranial Theta Burst Stimulation

We delivered direct electrical stimulation to the basolateral amygdala during half of the trials in the encoding phase of each experimental session. Stimulation pulses were delivered immediately once the image was removed from the screen and in a patterned rhythm designed to entrain endogenous theta-gamma oscillatory interactions (i.e., theta-burst stimulation, TBS) (Hanslmayr et al., 2016; Suppa et al., 2016). Specifically, we administered current-controlled, charge-balanced, bipolar, 1 mA, biphasic rectangular pulses over a 1 s period with a 50% duty cycle. Stimulation pulses were delivered at a rate of 50 Hz and nested within eight equally-spaced bursts (∼8 Hz) (*Figure 2B*–*C*). A subset of experiments (n = 7) used a lower current (0.5 mA) with variable pulse frequencies across trials (33 Hz, 50 Hz, 80 Hz).

### Peri-Stimulation Modulation Analyses

We first created peri-stimulation epochs (1 s pre-trial ISI, 1 s after image onset, 1 s during stimulation/after image offset, 1 s post-stimulation), with t = 0 representing stimulation onset and the moment at which the image was removed from the screen (*Figure 2A*); identical epochs were created for the image-only (no-stimulation) trials. Each epoch was constrained to 1 s to ensure that subsequent firing rate contrasts were unbiased and to capture potential transient effects (e.g., image onset/offset). Units with a trial-averaged baseline (pre-trial ISI) firing rate of < 0.1 Hz were excluded from subsequent analyses because low firing rates limited the ability to detect modulation robustly. Units were designated as “modulated” if either the during- or post-stimulation firing rate contrast was significant following permutation testing (described in *Statistical Analyses*). An additional contrast of pre-trial ISI vs. image onset was performed to evaluate the sensitivity of neurons to task stimuli (i.e., image presentation).

To investigate whether stimulation altered rhythmicity in neuronal firing, we analyzed the spike timing autocorrelograms. More specifically, we computed the pairwise differences in spike timing for each trial (bin size = 5 ms, max lag = 250 ms) and then contrasted the differences in the latencies of the peak normalized autocorrelation value between epochs (pre-, during-, post-stimulation). Only neurons with a firing rate of ≥ 1 Hz (n = 70/203, 34.5%) were included in this analysis since sparse firing resulted in noisy autocorrelation estimates.

Finally, we measured modulation in multiunit activity (MUA) by filtering the microleectrode signals in a 300-3,000 Hz window and counting the number of threshold crossings. Thresholds were determined on a per-channel basis and defined as -3.5 times the root mean square of the signal during the baseline period; activity during stimulation was excluded since stimulation artifact is difficult to separate from MUA in the absence of spike sorting.

### Population Analyses

To analyze pseudo-population activity, we performed a linear dimensionality reduction with PCA on a matrix of the z-scored trial-averaged firing rates of all neurons recorded across patients (*sklearn.decomposition.PCA*). This approach enables the identification of dominant patterns of coordinated neural activity that may not be apparent when examining individual neurons in isolation. Doing so allowed us to qualitatively examine the temporal dynamics of the dominant modes of neuron coactivity in a low-dimensional subspace, separated by experimental condition (stim vs. no-stim), region, and laterality relative to stimulation. The firing rate matrices for each condition were concatenated prior to PCA to facilitate direct comparison among the principal components; for simplicity, only the first three principal components are visualized. By collapsing across subjects into a common pseudo-population, this analysis provides a mesoscale view of how stimulation modulates shared activity patterns across anatomically distributed neural populations.

### Statistical Approach

All statistical analyses were conducted using custom Python scripts and established statistical libraries (i.e., Scikit-learn (Pedregosa et al., 2012), *Scipy* (Virtanen et al., 2020)*, Statsmodels* (Seabold and Perktold, 2010)). We performed two separate Wilcoxon signed-rank tests across trials on the during- and post-stimulation spike counts relative to their corresponding pre-trail baseline spike counts. To control for false positives, we compared the empirical test statistic against a null distribution generated from shuffling pre/during/post epoch labels (n = 1000 permutations) (*Figure 2B*). An identical analysis was also performed on the no-stimulation (image-only) trials.

To test for differences in the proportion of modulated units (across conditions, regions, cell type, stimulation parameters, and behavioral outcomes), we performed a series of one- and two-sided Fisher’s exact tests. Next, we used Mann-Whitney U tests to contrast baseline firing rates among modulated vs. unaffected units. Behavioral performance during the memory task was calculated using d-prime (d’), defined as the difference in an individual’s z-scored hit rate and false alarm rate. Observed changes in recognition memory were split into two categories using a d’ difference threshold of ± 0.5: responder (Δd’ < -.5 or Δd’ > +.5) or non-responder (-0.5 ≤ Δd’ ≤ 0.5). The threshold of ± 0.5 was chosen based on the defined range of a “medium effect” for Cohen’s *d,* which bears conceptual similarity to d’. To test the hypothesis that stimulation affected behavioral performance, we used a linear mixed effects model with d’ score as the dependent variable, condition and experiment as fixed effects, and session as a random effect; an additional test for differences among hit rates (percent of previously seen images correctly identified) was implemented using a paired-samples t-test.

### Firing Rate Control Analyses

We performed a series of control analyses to test whether our approach to firing rate detection was robust. First, we performed a sensitivity analysis by systematically varying the baseline firing rate threshold used to exclude units from modulation analyses. The threshold for inclusion of units was varied from 0–3 Hz (0.1 Hz step size), and the firing rate analyses were repeated to quantify the proportion of units meeting inclusion criteria and the proportion of units designated as modulated (*Figure S7A*). Next, we performed a dropout analysis wherein segments of data near the onset of a stimulation burst were removed from the during-stimulation epoch (an identical segment was also removed from the pre-trial ISI and post-stimulation epochs). To this end, we removed a window of data starting at the onset of each burst spanning 0–60 ms (5 ms step size, eight bursts in train) and recomputed the proportion of units meeting inclusion criteria and the proportion of units designated as modulated (*Figure S7B*). Finally, to better understand the tradeoffs with our statistical approach, we generated synthetic data with different baseline firing rates (0.1-5.0 Hz) and effect sizes (± 0.1-0.7 Hz), then simulated the likelihood of detecting modulation across variable conditions (*Figure S7C*).

## Results

We recorded single-unit activity from 23 patients (n = 30 sessions) with medically refractory epilepsy as they completed a visual recognition memory task (see *Table S1* for patient demographics). During the encoding session of the experiment, each patient received either 80 or 160 trials of bipolar intracranial TBS to a contiguous pair of macroelectrode contacts in the BLA (see *Figure S1* for anatomical localization of stimulated contacts. An equal number of “no-stimulation” trials were randomly interspersed to evaluate the effect of stimulation on memory performance and control for neuronal modulation resulting from experimental stimuli (e.g., image presentation). In total, we isolated 203 putative neurons from 68 bundles of 8 microwires each, distributed among recording sites in the hippocampus (HIP, n = 95 units), orbitofrontal cortex (OFC, n = 44), amygdala (AMY, n = 39), and anterior cingulate cortex (ACC, n = 25) (*Figure 1*; see also *Figure S2* and *Figure S3*); a subset of these units (n = 47, 23.2%) was excluded from subsequent analyses because low baseline firing rates (< 0.1 Hz) limited the ability to robustly detect modulation.

### Theta Burst Stimulation of the BLA Modulates Widely Distributed Populations of Neurons

We hypothesized that BLA stimulation would modulate neuronal activity in the sampled regions, given the amygdala’s well-established connectivity to the HIP, OFC, and ACC (Roy et al., 2009). To test this hypothesis, we quantified spike counts across trials within peri-stimulation epochs (1 s pre-trial ISI, 1 s after image onset, 1 s during stimulation/after image offset, and 1 s post-stimulation) and used Wilcoxon signed-rank tests to compare the spike counts against a null distribution generated by shuffling epoch labels. We performed two firing rate contrasts across trials (pre-trial ISI vs. during stimulation, pre-trial ISI vs. post-stimulation) within two distinct conditions (stim, no-stim); an additional contrast of the pre-trial ISI vs. image onset epochs was included to evaluate the sensitivity of neurons to task image presentations (*Figure 2A*).

BLA TBS modulated firing rates in 30.1% of all recorded units, a significantly higher proportion than the 15.4% responsive to no-stim (image only) trials (one-sided Fisher’s exact test, OR = 2.37, *p* < 0.001; *Figure 3A*). Across all regions sampled, we observed units modulated by the stim and no-stim conditions. Units in HIP (OR = 2.07, *p* = 0.044), OFC (OR = 5.09, *p* = 0.040), and AMY (OR = 3.33, *p* = 0.042) were most sensitive to stimulation; we did not observe a difference in the proportions of units within the ACC responsive to the stim vs. no-stim conditions (OR = 1.00, *p* = 0.661; *Figure 3B*). Only 9.0% of units responded to both the stim and no-stim conditions, despite approximately half of the stim-modulated units (representing 14.7% of all units) exhibiting a change in firing rate associated with image onset (*Figure 3D*). This result suggests that the units modulated by stimulation are largely distinct from those responsive to image offset during trials without stimulation. Stimulation, however, did not appear to alter the rhythmicity in neuronal firing, as measured by spiking autocorrelograms (*Figure S5*).

Because neuronal firing properties vary across cell types (Barthó et al., 2004; Keller et al., 2010; Peyrache et al., 2012; Quyen et al., 2008), we also tested whether baseline (pre-trial ISI) firing rates predicted a unit’s response to stimulation, suggestive of selective engagement of specific neuron populations. Stimulation-modulated units exhibited significantly higher baseline firing rates compared to unaffected units (*U*(N_Stim, Mod_ = 47, N_Stim, NS_ = 109) = 3450.50, *p* < 0.001). No difference in baseline firing rate was observed among units modulated in the no-stim condition, compared to those that were unaffected (*U*(N_No-Stim, Mod_ = 24, N_No-Stim, NS_ = 132) = 1964.00, *p* = 0.062) (*Figure 3C*). The median (Q1, Q3) baseline firing rates for modulated units in the stim and no-stim conditions were 1.77 Hz (0.95 Hz, 5.39 Hz) and 1.53 Hz (0.72 Hz, 5.41 Hz), respectively.

In a subset of experimental sessions (n = 7), we explored the effects of different stimulation parameters on neuronal modulation within an experimental session; more specifically, we employed a lower stimulation amplitude (0.5 mA vs. 1.0 mA) and varied pulse frequency (33 Hz vs. 50 Hz and 33 Hz vs. 80 Hz). Neither the amplitude of stimulation (OR = 1.69, *p* = 0.302, n = 30) nor pulse frequency (33 vs. 80 Hz; OR = 0.00, *p* = 1.000, n = 1; 50 vs. 80 Hz; OR = 1.40, *p* = 0.758, n = 6) significantly altered the proportion of modulated units. (*Figure S4B*).

Our exploratory analyses of pseudo-population activity revealed interesting temporal dynamics associated with image presentation and the delivery of stimulation. More specifically, we observed variation among the first three principal components across both stim and no-stim trials associated with image presentation (t = -3 to t = 0) and robust shifts in coactivity modes associated with stimulation (t = 0 to t = 1) (*Figure 3F*). Further characterization of these dynamics suggests that TBS of the BLA primarily resulted in transient changes to firing rate coactivity within hippocampal neurons and was present regardless of the neuron’s laterality to stimulation (*Figure S6A–B*). Taken together, these analyses reveal global structure in the state space of responses to BLA stimulation within hippocampal circuits.

Finally, we performed three supplementary analyses to evaluate the robustness of our approach to detecting firing rate modulation: a sensitivity analysis assessing the proportion of modulated units at different firing rate thresholds for inclusion/exclusion, a data dropout analysis designed to control for the possibility that non-physiological stimulation artifacts may preclude the detection of temporally adjacent spiking, and a synthetic detection probability analysis. These results recapitulate our observation that units with higher baseline firing are most likely to exhibit modulation (though the probability of detecting modulation is lower for sparsely active neurons) and suggest that suppression in firing rate is not solely attributable to amplifier saturation following stimulation (*Figure S7*).

### Neurons Exhibit Heterogenous Responses to Theta Burst Stimulation

Recent studies have reported enhanced neural plasticity (via intracranial local field potential recordings and evoked responses) following repetitive direct electrical stimulation (Herrero et al., 2021; Huang et al., 2024, 2019; Keller et al., 2018). Accordingly, we hypothesized that recorded units would predominantly exhibit enhanced spiking in response to intracranial TBS of the BLA. Similarly, individual units exhibited highly variable responses to stimulation with respect to onset latency (rapid vs. delayed), duration (transient vs. durable), and valence (enhancement vs. suppression) (*Figure 2B*).

The most common epoch for firing rate modulation was during the 1 s epoch in which TBS was delivered (25.0% of all neurons). Smaller subsets were modulated only in the 1 s post-stimulation epoch (6.4%) or in both the during- and post-stimulation epochs (1.3%). A similar trend was observed for modulation in the no-stim condition: 10.9% during, 5.8% post, and 1.3% for both. Suppression was most common among modulated units during stimulation (56.4%), whereas enhancement was the dominant response post-stimulation (70.0%). In contrast, enhancement was most common within both epochs across no-stim trials (58.5% during, 66.7% post). The mean (± SD) absolute z-scored difference in firing rate across stimulation trials (relative to pre-trial ISI) was z = 0.60 (± 0.58) and z = 0.43 (± 0.27) for the during- and post-stimulation epochs, respectively. Across no-stim trials, we observed a mean absolute z-scored difference of z = 0.38 (± 0.24) and z = 0.30 (± 0.18) in analogous epochs (*Figure 3E*). Additional characterization of MUA revealed a dominant signature of increased activity post-vs. pre-stimulation, in line with these trends observed at the single-neuron level (*Figure S8*).

### Theta Burst Stimulation Modulates Excitatory and Inhibitory Neurons Equally

Using k-means clustering, we grouped neurons into two distinct clusters based on waveform morphology, representing neurons that were presumed to be excitatory (E) and inhibitory (I) (*Figure 4B*). Inhibitory (fast-spiking) neurons exhibited shorter waveform VP and PHW, compared with excitatory (regular-spiking) neurons (I cluster centroid: VP = 0.50ms, PHW = 0.51ms; E cluster centroid: VP = 0.32ms, PHW = 0.31ms), and greater baseline firing rates (*U*(N_I_ = 23, N_E_ = 133) = 1074.50, *p* = 0.023) (*Figure 4D*). Although we observed a much greater proportion of excitatory vs. inhibitory neurons (E: 85.3%, I: 14.7%), stimulation appeared to affect excitatory and inhibitory neurons equally, suggesting that one cell type is not preferentially activated over another (*Figure 4E*).

### Association Between Neuronal Modulation and Memory Performance is Unclear

Next, we investigated the link between stimulation and performance during the visual recognition memory task. To this end, we first used a linear mixed-effects model to examine the effect of condition (stim, no-stim) on memory performance (d’) across trials in each session, with individual sessions treated as a random effect (intercept). Experiment type was also included as a fixed effect since data were aggregated across four highly similar experiments with minor differences in the content of visual stimuli, number of trials, stimulation parameters, and testing intervals. We did not observe an overall effect of memory enhancement (*p* > 0.05) when controlling for subject-level variability (*Figure 5A*). The lack of a memory enhancement effect may be associated with high hit rates limiting sensitivity (mean ± SD) (75.7% ± 13.5% for no-stim trials, 75.0% ± 14.3% for stim trials), and considerable variability among false alarm rates across participants (17.9% ± 17.4%, range 0–70%; *Figure 5B*).

**Figure 5:**
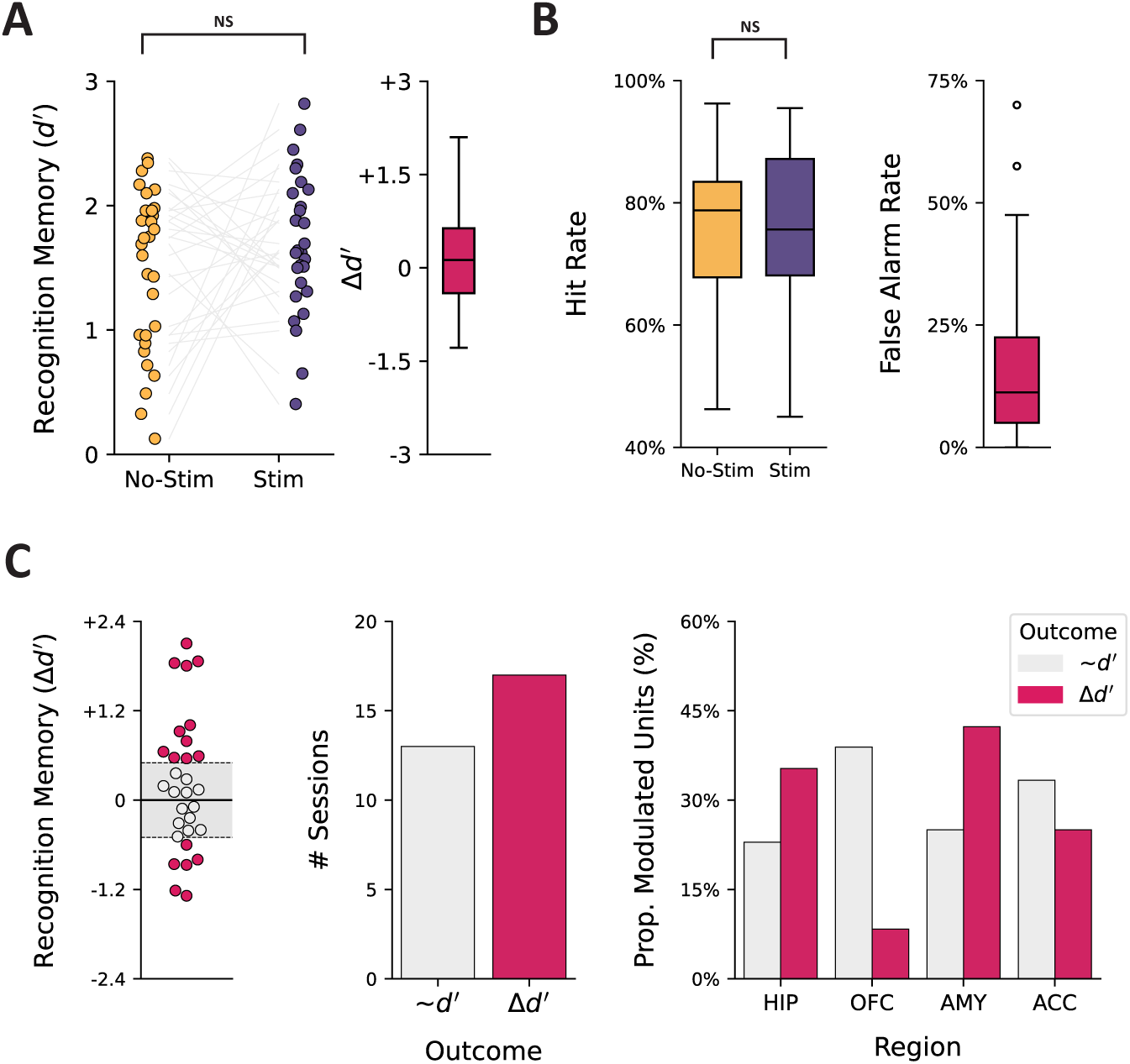
Summary of behavioral performance during memory task. (A) Memory performance for each session is quantified using d’ (left); grey lines connect d’ scores across conditions for an individual session. Boxplot of the observed difference in d’ scores across conditions (right). (B) Hit rate (percent of old images correctly recognized) and false alarm rate (percent of new images incorrectly labeled as old) across conditions. (C) Change in recognition memory performance was split into two categories using a d’ difference threshold of ± 0.5: responder (positive or negative; Δd’, pink) and non-responder (∼d’, grey). Individual d’ scores are shown (left) with points colored by outcome category; dotted lines demarcate category boundaries, and the grey-shaded region represents negligible change. The number of sessions within each outcome category (middle) and the proportion of modulated units as a function of outcome category, separated by region (right). NS = not significant.

At the level of individual sessions, we observed enhanced memory (Δd’ > +0.5) in 36.7%, impaired memory (Δd’ < -0.5) in 20.0%, and negligible change (-0.5 ≤ Δd’ ≤ 0.5) in 43.3% when comparing performance between the stim and no-stim conditions; a threshold of Δd’ ± 0.5 was chosen for this classification based on the defined range of a “medium effect” for Cohen’s *d*. To test our hypothesis that neuronal modulation would be associated with changes in memory performance, we combined the sessions that resulted in either memory enhancement or impairment and contrasted the proportion of modulated units across regions sampled. We did not, however, observe a meaningful difference in the proportion of modulated units when grouped by behavioral outcome (all contrasts *p* > 0.05) (*Figure 5C*).

Proximity to white matter has been shown to influence the effects of stimulation on behavior and the strength of evoked responses (Mankin et al., 2021; Mohan et al., 2020; Paulk et al., 2022). Across all stimulated contacts, we observed only small differences in the proximity of stimulation contacts to white matter (median = 4.6 mm, range = 1.5-8.0 mm), likely because the chosen target (i.e., basolateral amygdala) has several nearby white matter structures (e.g., stria terminalis). Nonetheless, we performed a linear regression between the proximity to white matter and the stimulation-induced effect on behavior (stimulation vs. no-stimulation *d’* difference), the results of which indicate no clear association (*p* > 0.05; see *Figure S9*).

## Discussion

Theta-burst stimulation is an efficient and validated paradigm for inducing long-term potentiation (LTP) in neural circuits (Larson and Munkácsy, 2015). Additionally, intracranial TBS was recently shown to promote region-specific short-term plasticity (Herrero et al., 2021; Huang et al., 2024) and entrain frequency-matched oscillations (Solomon et al., 2021). At present, however, there is an incomplete understanding of how these population-level responses to stimulation relate to a modulation in the activity of individual neurons, which are thought to be the substrate of memory encoding and retrieval (Hebb, 1949). Here, we address this knowledge gap by characterizing neuronal firing recorded from microelectrodes in humans undergoing intracranial TBS of the BLA. Our experimental design focused explicitly on stimulation of the BLA, given our prior work that seeks to understand amygdala-mediated memory enhancement in humans (Inman et al., 2023, 2020, 2018).

We observed neurons distributed throughout the hippocampus, orbitofrontal cortex, anterior cingulate cortex, and amygdala responsive to direct electrical TBS. The effect of TBS on firing rate was heterogeneous with respect to onset, duration, and valence. Previous work characterizing local-field potential responses to intracranial TBS observed similarly bidirectional effects throughout the brain suggestive of short-term plasticity (Huang et al., 2024). Few studies, however, have characterized the impact of exogenous stimulation on the spiking activity of individual neurons, none of which involve intracranial theta burst stimulation. One study reported a long-lasting reduction in neural excitability among parietal neurons, with variable onset time and recovery following continuous transcranial TBS in non-human primates (Romero et al., 2022). In a similar vein, it was recently shown that human neurons are largely suppressed by single-pulse electrical stimulation (Cowan et al., 2024; Plas et al., 2024). Other emerging evidence suggests that transcranial direct current stimulation may entrain the rhythm rather than rate of neuronal spiking (Krause et al., 2019) and that stimulation-evoked modulation of spiking may meaningfully impact behavioral performance on cognitive tasks (Fehring et al., 2024). An alternative approach has focused on the delivery of spatially selective microstimulation resembling the extracellular currents that normally modulate neuronal activity—this methodology has been used to bi-directionally drive neuronal firing in human temporal cortex (Youssef et al., 2023) and enhance memory specificity for images following stimulation (Titiz et al., 2017).

Although subsets of neurons from each region we sampled were responsive to stimulation, we observed the greatest difference in the proportion of modulated units across conditions in the hippocampus, orbitofrontal cortex, and amygdala. This regional selectivity is to be expected, given that numerous studies have characterized how structural, functional, and effective connectivity among brain regions predicts the effects of stimulation (Fox et al., 2020; Huang et al., 2024, 2019; Keller et al., 2018, 2011; Solomon et al., 2018; Stiso et al., 2019). We also observed that units with greater baseline activity were most likely to exhibit modulated firing rates following stimulation. Other studies have identified firing patterns and waveform properties that differ between inhibitory and excitatory neurons in humans (Barthó et al., 2004; Keller et al., 2010; Peyrache et al., 2012; Quyen et al., 2008). For example, baseline firing rate disambiguates “regular-spiking” and “fast-spiking” neurons, which are presumed to represent pyramidal cells and interneurons, respectively. To test these hypotheses directly, we clustered neurons into presumed excitatory and inhibitory neurons based on waveform morphology. In doing so, we observed ∼85% excitatory and ∼15% inhibitory neurons, which is very similar to what has been reported previously in human intracranial recordings (Cowan et al., 2024; Peyrache et al., 2012). Interestingly, stimulation appeared to modulate approximately the same proportion of neurons for each cell type (∼30%), despite the differently-sized groups. Recent reports, however, have suggested that the extent to which electrical fields entrain neuronal spiking, particularly with respect to phase-locking, may be specific to distinct classes of cells (Lee et al., 2024).

Modulation in neuronal activity was defined by contrasting firing rates before, during, and after TBS across trials. In doing so, we were able to characterize coarse differences in activity indicative of enhancement or suppression. This approach, however, did not allow for analysis of more subtle, nuanced effects such as entrainment of spiking to individual bursts or pulses of TBS. Characterizations of rhythmicity in firing were challenging, given that most of the neurons we identified exhibit sparse activity with low baseline firing rates, and stimulation often resulted in further suppression of spiking.

Although stimulation artifacts resulted in amplitude threshold crossings that may be spuriously interpreted as a neuronal spike, we implemented several methods to mitigate the influence of non-physiological activity. First, the characteristics of each unit (e.g., waveform shape) were manually inspected during spike sorting and further quantified using several quality control metrics (e.g., interspike intervals); stimulation resulted in a stereotyped response that was easily detectable and removed from subsequent analyses. Additionally, we tested for modulation both during stimulation and in the post-stimulation epochs—a period in which no artifact was present. Contrary to what would be expected if stimulation artifacts were falsely elevating firing rates, we observed predominantly suppression during stimulation and enhancement post-stimulation.

Since we collected our microelectrode recordings in the context of a visual recognition memory task, we tested whether stimulation resulted in a change in memory performance. Although we hypothesized that stimulation would improve memory performance, we did not identify an apparent stimulation-related memory enhancement when controlling for individual differences, in contrast to our prior work (Inman et al., 2018). The absence of such an effect may be, at least in part, attributable to the considerable variability that we observed in this cohort; indeed, baseline memory performance and individual differences have been shown to account for a substantial portion of the variability in this amygdala-mediated memory enhancement effect (Hollearn et al., 2024).

Several studies on rats have demonstrated that brief electrical stimulation of the BLA can prioritize the consolidation of specific memories (Bass et al., 2014, 2012; Bass and Manns, 2015; Manns and Bass, 2016). These pro-memory effects emerged ∼24 hours post-encoding and appear to be hippocampal-dependent (Bass et al., 2014), despite not resulting in a net change in the firing rates of hippocampal pyramidal neurons; instead, BLA stimulation resulted in brief periods of spike-field and field-field synchrony within CA3–CA1 in the low-gamma frequency range (30–55 Hz), which may facilitate spike-timing-dependent plasticity in recently active neurons (Bass and Manns, 2015).

The present study did not investigate interactions between spiking activity and local field potentials because neuronal spiking was sparse at baseline and often further suppressed by stimulation; only a very small proportion of the total number of trials across all neurons exhibited ≥ 10 spikes in both the 1 s pre- and post-stimulation epochs (∼2.5%). Although certain metrics are not biased by sample size (e.g., pairwise phase consistency), low spike counts can dramatically affect variance and, therefore, result in unstable estimates (Vinck et al., 2011).

How exactly the activity of single neurons is aggregated to produce local field potentials, which in turn interact with neuronal ensembles distributed throughout the brain, remains an active area of research (Herreras, 2016; Kajikawa and Schroeder, 2011; Manning et al., 2009; Teleńczuk et al., 2020). One recent study that leveraged closed-loop stimulation targeting memory consolidation during sleep observed neuronal spiking with greater phase-locking to medial temporal lobe slow-wave activity following stimulation (Geva-Sagiv et al., 2023); neuronal phase-locking, particularly to hippocampal theta oscillations, has long been associated with robust memory encoding and retrieval (Jacobs et al., 2007; Rutishauser et al., 2010; Schonhaut et al., 2024; Yoo et al., 2021). Further characterization of these spike-field interactions and refinement of closed-loop stimulation methods may provide a means for precisely modulating neuronal dynamics, for example, by entraining neuronal spiking that is phase-aligned to endogenous hippocampal theta oscillations to selectively enhance the encoding or retrieval of memories (Hasselmo, 2005; Hasselmo et al., 2002; Hasselmo and Stern, 2014; Siegle and Wilson, 2014).

Finally, we performed an exploratory analysis of neuronal pseudo-population activity, which suggests that hippocampal neurons exhibit robust changes in firing rate coactivity in response to BLA stimulation. Related research has similarly described how BLA stimulation can induce synchronous firing of hippocampal neurons, which has memory-enhancing effects (Bass and Manns, 2015). Other studies have leveraged similar, low-dimensional representations of population dynamics to describe how coordinated neural activity facilitates inferential reasoning and memory retrieval within the medial frontal cortex and hippocampus, respectively (Courellis et al., 2024; Minxha et al., 2020). Thus, a greater understanding of how neuronal coactivity may be precisely modulated by stimulation may help to refine therapeutic interventions targeting complex cognition and computation.

## Conclusions

By characterizing patterns of neuronal modulation evoked by intracranial TBS, we provide new insights that link micro- and macroscale responses to stimulation of the human brain. These insights advance our limited understanding of how focal electrical fields influence neuronal firing at the single-cell level and motivate future neuromodulatory therapies that aim to recapitulate specific patterns of activity implicated in cognition and memory.

## CRediT Author Statement

**Justin Campbell**: Conceptualization, Methodology, Software, Formal Analysis, Investigation, Data Curation, Writing – Original Draft, Writing – Review & Editing, Visualization; **Rhiannon Cowan**: Conceptualization, Data Curation, Writing – Review & Editing; **Krista Wahlstrom**: Investigation, Writing – Review & Editing, Project Administration; **Martina Hollearn**: Writing – Review & Editing; **Dylan Jensen**: Writing – Review & Editing; **Tyler Davis**: Software, Writing – Review & Editing; **Shervin Rahimpour**: Resources, Writing – Review & Editing; **Ben Shofty**: Resources, Writing – Review & Editing; **Amir Arain**: Resources, Writing – Review & Editing; **John Rolston**: Resources, Writing – Review & Editing; **Stephan Hamann**: Writing – Review & Editing; **Shuo Wang**: Resources, Writing – Review & Editing; **Lawrence Eisenman**: Resources, Writing – Review & Editing; **James Swift**: Software, Investigation, Writing – Review & Editing; **Tao Xie**: Investigation, Writing – Review & Editing; **Peter Brunner**: Software, Investigation, Supervision, Writing – Review & Editing; **Joe Manns**: Conceptualization, Supervision, Project Administration, Writing – Review & Editing; **Cory Inman**: Supervision, Resources, Project Administration, Funding Acquisition, Writing – Review & Editing; **Elliot Smith**: Conceptualization, Supervision, Resources, Writing – Review & Editing; **Jon Willie**: Investigation, Supervision, Resources, Project Administration, Funding Acquisition, Writing – Review & Editing.

## Resource Availability

Custom Python analysis scripts used in the manuscript are publicly available on GitHub (https://github.com/Justin-Campbell/BLAESUnits). Deidentified neural recordings may be made available upon reasonable request.

## Acknowledgments

We are grateful to all the patients who participated in the study.

---

**Supplemental Figure 1:** Anatomical location of stimulated electrodes.

A coronal slice from the T1-weighted MRI scan is shown for each patient who participated in the study (n = 16). Electrode contacts within the same plane of the image are shown with blue circles, and the bipolar pair of stimulated contacts within the basolateral amygdala is highlighted in red.

**Supplemental Figure 2: Unit quality metrics.**

(A) Number of units detected per implanted microelectrode bundle. (B) Mean firing rate (Hz) across recording session. (C) Percent of interspike intervals < 3 ms. (D) Interspike interval coefficient of variation. (E) Mean presence ratio of firing within units (1 s bins). (F) Signal-to-noise ratio of unit waveform peak. (G) Mean signal-to-noise ratio across the entire unit waveform. (H) Representative example of stereotyped, high-amplitude stimulation-artifact waveform; non-physiological waveforms were excluded from analysis.

**Supplemental Figure 3 Characterization of units based on laterality relative to stimulation.**

(A) Unit counts on the contralateral (Contra) or ipsilateral (Ipsi) side of stimulation. (B) Unit counts separated by laterality and region. (C) Stacked histograms of Euclidean distance between microelectrode bundle location and stimulation contacts, separated by laterality (bin size = 5 mm).

**Supplemental Figure 4 Sub-analysis of stimulation parameters used across experiments.**

Comparison of the proportion of stimulation-modulated units across sessions with 1 mA (n = 23) vs. 0.5 mA (n = 7). (B) Comparison of the proportion of stimulation-modulated units across sessions testing distinct pulse frequencies: 33 Hz vs. 80 Hz (n = 1) and 50 Hz vs. 80 Hz (n = 6). The values above individual bars represent the number of sessions using that stimulation parameter. NS = not significant.

**Supplemental Figure 5 Analysis of modulation in spiking rhythmicity.**

(A) Representative autocorrelograms ACG) for a single neuron. The pairwise differences in spike timing were computed for each trial and epoch (bin size = 5 ms, max lag = 250 ms), then smoothed with a Gaussian kernel. The peak in the normalized ACG across trials was computed for each epoch. (B) Kernel density estimate of the peak ACG lag, separated by epoch. (C) The peak ACG lags were split by whether the neuron was modulated (Mod) or unaffected by stimulation (NS = not significant) for each of the two contrasts: pre-vs. during-stim (left) and pre-vs. post-stim (right).

**Supplemental Figure 6 Analysis of pseudo-population activity within regions, separated by laterality relative to stimulation.**

Pseudo-population activity was characterized within each region via a linear dimensionality reduction on the trial-averaged firing rates. The temporal dynamics of each region’s first three principal components are shown for (A) units ipsilateral to stimulation and (B) units contralateral to stimulation. HIP = hippocampus (coral), OFC = orbitofrontal cortex (yellow), AMY = amygdala (blue), ACC = anterior cingulate cortex (purple).

**Supplemental Figure 7: Control analyses for the detection of modulated units.**

The same permutation-based analyses reported in the manuscript were repeated under different control conditions. The percent of units (total n = 203) that met the firing rate threshold for inclusion (pink), the percent of included units modulated in the stim condition (purple), and the percent of included units modulated in the no-stim condition are shown. (A) The threshold for inclusion of units was varied from 0–3 Hz (0.1 step size); the black dashed line represents the ≥ 0.1 Hz threshold used in the manuscript. (B) To control for the possibility that non-physiological stimulation artifacts may preclude the detection of temporally adjacent spiking, we removed segments of data beginning at the onset of each burst of pulses (0–60 ms, 5 ms step size). Identical temporal windows were removed from the corresponding pre- and post-stimulation epochs to mitigate effects resulting solely from summation over different epoch sizes (reduced spike counts with shorter windows). (C) Visualization of the predicted probability of detecting modulation across synthetic neurons with variable firing rates and modulation effect sizes; FR = firing rate.

**Supplemental Figure 8: Analysis of multiunit activity response to stimulation.**

(A) Example trace of multiunit activity (MUA) in one channel during a single stimulation trial. Threshold crossings are highlighted with a pink dot overlaid on the MUA signal with a corresponding hash below. (B) The percentage of channels with significantly modulated MUA, separated by the direction of effect. (C) The percentage of channels with significantly modulated MUA, separated by direction effect and region. Inc (red; post > pre) vs. Dec (blue; post < pre). HIP = hippocampus, OFC = orbitofrontal cortex, AMY = amygdala, ACC = anterior cingulate cortex. *** *p* < 0.001, NS = not significant.

**Supplemental Figure 9: The effect of stimulation proximity to white matter and distance to recorded neurons**

(A) Kernel density estimate of the Euclidean distance from stimulation contacts to nearest WM structure (in mm); hash marks represent individual observations. (B) The change in memory performance (Δd’) was linearly regressed onto the distance from the stimulated contacts to white matter.

## References

1. Barthó P, Hirase H, Monconduit L, Zugaro M, Harris KD, Buzsáki G. 2004. Characterization of Neocortical Principal Cells and Interneurons by Network Interactions and Extracellular Features. J Neurophysiol 92:600–608. doi:10.1152/jn.01170.2003

2. Bass DI, Manns JR. 2015. Memory-Enhancing Amygdala Stimulation Elicits Gamma Synchrony in the Hippocampus. Behav Neurosci 129:244–256. doi:10.1037/bne0000052

3. Bass DI, Nizam ZG, Partain KN, Wang A, Manns JR. 2014. Amygdala-mediated enhancement of memory for specific events depends on the hippocampus. Neurobiol Learn Mem 107:37–41. doi:10.1016/j.nlm.2013.10.020

4. Bass DI, Partain KN, Manns JR. 2012. Event-Specific Enhancement of Memory via Brief Electrical Stimulation to the Basolateral Complex of the Amygdala in Rats. Behav Neurosci 126:204–208. doi:10.1037/a0026462

5. Courellis HS, Minxha J, Cardenas AR, Kimmel DL, Reed CM, Valiante TA, Salzman CD, Mamelak AN, Fusi S, Rutishauser U. 2024. Abstract representations emerge in human hippocampal neurons during inference. Nature 1–9. doi:10.1038/s41586-024-07799-x

6. Cowan RL, Davis TS, Merricks EM, Kundu B, Shofty B, Rahimpour S, Schevon CA, Rolston JD, Smith EH. 2024. Cell-type specific responses to single-pulse electrical stimulation of the human brain. bioRxiv 2024.11.24.625077. doi:10.1101/2024.11.24.625077

7. Davis TS, Caston RM, Philip B, Charlebois CM, Anderson DN, Weaver KE, Smith EH, Rolston JD. 2021. LeGUI: A Fast and Accurate Graphical User Interface for Automated Detection and Anatomical Localization of Intracranial Electrodes. Front Neurosci 15:769872. doi:10.3389/fnins.2021.769872

8. Dolcos F, LaBar KS, Cabeza R. 2004. Interaction between the Amygdala and the Medial Temporal Lobe Memory System Predicts Better Memory for Emotional Events. Neuron 42:855–863. doi:10.1016/s0896-6273(04)00289-2

9. Fehring DJ, Yokoo S, Abe H, Buckley MJ, Miyamoto K, Jaberzadeh S, Yamamori T, Tanaka K, Rosa MGP, Mansouri FA. 2024. Direct current stimulation modulates prefrontal cell activity and behaviour without inducing seizure-like firing. Brain awae273. doi:10.1093/brain/awae273

10. Fox KCR, Shi L, Baek S, Raccah O, Foster BL, Saha S, Margulies DS, Kucyi A, Parvizi J. 2020. Intrinsic network architecture predicts the effects elicited by intracranial electrical stimulation of the human brain. Nat Hum Behav 4:1039–1052. doi:10.1038/s41562-020-0910-1

11. Geva-Sagiv M, Mankin EA, Eliashiv D, Epstein S, Cherry N, Kalender G, Tchemodanov N, Nir Y, Fried I. 2023. Augmenting hippocampal–prefrontal neuronal synchrony during sleep enhances memory consolidation in humans. Nat Neurosci 1–11. doi:10.1038/s41593-023-01324-5

12. Hanslmayr S, Staresina BP, Bowman H. 2016. Oscillations and Episodic Memory: Addressing the Synchronization/Desynchronization Conundrum. Trends Neurosci 39:16–25. doi:10.1016/j.tins.2015.11.004

13. Hasselmo ME. 2005. What is the function of hippocampal theta rhythm?—Linking behavioral data to phasic properties of field potential and unit recording data. Hippocampus 15:936–949. doi:10.1002/hipo.20116

14. Hasselmo ME, Bodeln C, Wyble BP. 2002. A Proposed Function for Hippocampal Theta Rhythm: Separate Phases of Encoding and Retrieval Enhance Reversal of Prior Learning. Neural Comput 14:793–817. doi:10.1162/089976602317318965

15. Hasselmo ME, Stern CE. 2014. Theta rhythm and the encoding and retrieval of space and time. Neuroimage 85:656–666. doi:10.1016/j.neuroimage.2013.06.022

16. Hebb DO. 1949. The organization of behavior; A neuropsychological theory. New York: John Wiley and Sons, Inc. doi:10.1002/sce.37303405110

17. Hermans EJ, Battaglia FP, Atsak P, Voogd LD de, Fernández G, Roozendaal B. 2014. How the amygdala affects emotional memory by altering brain network properties. Neurobiol Learn Mem 112:2–16. doi:10.1016/j.nlm.2014.02.005

18. Herreras O. 2016. Local Field Potentials: Myths and Misunderstandings. Front Neural Circuits 10:101. doi:10.3389/fncir.2016.00101

19. Herrero JL, Smith A, Mishra A, Markowitz N, Mehta AD, Bickel S. 2021. Inducing neuroplasticity through intracranial θ-burst stimulation in the human sensorimotor cortex. J Neurophysiol 126:1723–1739. doi:10.1152/jn.00320.2021

20. Hollearn MK, Manns JR, Blanpain LT, Hamann SB, Bijanki K, Gross RE, Drane DL, Campbell JM, Wahlstrom KL, Light GF, Tasevac A, Demarest P, Brunner P, Willie JT, Inman CS. 2024. Exploring individual differences in amygdala-mediated memory modulation. *Cogn, Affect*, Behav Neurosci 1–22. doi:10.3758/s13415-024-01250-4

21. Huang Y, Hajnal B, Entz L, Fabó D, Herrero JL, Mehta AD, Keller CJ. 2019. Intracortical Dynamics Underlying Repetitive Stimulation Predicts Changes in Network Connectivity. J Neurosci 39:6122–6135. doi:10.1523/jneurosci.0535-19.2019

22. Huang Y, Zelmann R, Hadar P, Dezha-Peralta J, Richardson RM, Williams ZM, Cash SS, Keller CJ, Paulk AC. 2024. Theta-burst direct electrical stimulation remodels human brain networks. Nat Commun 15:6982. doi:10.1038/s41467-024-51443-1

23. Inman CS, Bijanki KR, Bass DI, Gross RE, Hamann S, Willie JT. 2020. Human amygdala stimulation effects on emotion physiology and emotional experience. Neuropsychologia 145:106722. doi:10.1016/j.neuropsychologia.2018.03.019

24. Inman CS, Hollearn MK, Augustin L, Campbell JM, Olson KL, Wahlstrom KL. 2023. Discovering how the amygdala shapes human behavior: From lesion studies to neuromodulation. Neuron. doi:10.1016/j.neuron.2023.09.040

25. Inman CS, Manns JR, Bijanki KR, Bass DI, Hamann S, Drane DL, Fasano RE, Kovach CK, Gross RE, Willie JT. 2018. Direct electrical stimulation of the amygdala enhances declarative memory in humans. Proc National Acad Sci 115:98–103. doi:10.1073/pnas.1714058114

26. Jacobs J, Kahana MJ, Ekstrom AD, Fried I. 2007. Brain Oscillations Control Timing of Single-Neuron Activity in Humans. J Neurosci 27:3839–3844. doi:10.1523/jneurosci.4636-06.2007

27. Kajikawa Y, Schroeder CE. 2011. How Local Is the Local Field Potential? Neuron 72:847–858. doi:10.1016/j.neuron.2011.09.029

28. Keller CJ, Bickel S, Entz L, Ulbert I, Milham MP, Kelly C, Mehta AD. 2011. Intrinsic functional architecture predicts electrically evoked responses in the human brain. Proc Natl Acad Sci 108:10308–10313. doi:10.1073/pnas.1019750108

29. Keller CJ, Huang Y, Herrero JL, Fini M, Du V, Lado FA, Honey CJ, Mehta AD. 2018. Induction and quantification of excitability changes in human cortical networks. J Neurosci 38:1088–17. doi:10.1523/jneurosci.1088-17.2018

30. Keller CJ, Truccolo W, Gale JT, Eskandar E, Thesen T, Carlson C, Devinsky O, Kuzniecky R, Doyle WK, Madsen JR, Schomer DL, Mehta AD, Brown EN, Hochberg LR, Ulbert I, Halgren E, Cash SS. 2010. Heterogeneous neuronal firing patterns during interictal epileptiform discharges in the human cortex. Brain 133:1668–1681. doi:10.1093/brain/awq112

31. Kim MJ, Loucks RA, Palmer AL, Brown AC, Solomon KM, Marchante AN, Whalen PJ. 2011. The structural and functional connectivity of the amygdala: From normal emotion to pathological anxiety. Behav Brain Res 223:403–410. doi:10.1016/j.bbr.2011.04.025

32. Krause MR, Vieira PG, Csorba BA, Pilly PK, Pack CC. 2019. Transcranial alternating current stimulation entrains single-neuron activity in the primate brain. Proc Natl Acad Sci 116:5747–5755. doi:10.1073/pnas.1815958116

33. LaBar KS, Cabeza R. 2006. Cognitive neuroscience of emotional memory. Nat Rev Neurosci 7:54–64. doi:10.1038/nrn1825

34. Larson J, Munkácsy E. 2015. Theta-burst LTP. Brain Res 1621:38–50. doi:10.1016/j.brainres.2014.10.034

35. Lee SY, Kozalakis K, Baftizadeh F, Campagnola L, Jarsky T, Koch C, Anastassiou CA. 2024. Cell-class-specific electric field entrainment of neural activity. Neuron. doi:10.1016/j.neuron.2024.05.009

36. Mankin EA, Aghajan ZM, Schuette P, Tran ME, Tchemodanov N, Titiz A, Kalender G, Eliashiv D, Stern J, Weiss SA, Kirsch D, Knowlton B, Fried I, Suthana N. 2021. Stimulation of the right entorhinal white matter enhances visual memory encoding in humans. Brain Stimul 14:131–140. doi:10.1016/j.brs.2020.11.015

37. Manning JR, Jacobs J, Fried I, Kahana MJ. 2009. Broadband Shifts in Local Field Potential Power Spectra Are Correlated with Single-Neuron Spiking in Humans. J Neurosci 29:13613–13620. doi:10.1523/jneurosci.2041-09.2009

38. Manns JR, Bass DI. 2016. The Amygdala and Prioritization of Declarative Memories. Curr Dir Psychol Sci 25:261–265. doi:10.1177/0963721416654456

39. McDonald AJ, Mott DD. 2017. Functional neuroanatomy of amygdalohippocampal interconnections and their role in learning and memory. J Neurosci Res 95:797–820. doi:10.1002/jnr.23709

40. McGaugh JL. 2013. Making lasting memories: Remembering the significant. Proc Natl Acad Sci 110:10402–10407. doi:10.1073/pnas.1301209110

41. McGaugh JL. 2004. The Amygdala Modulates The Consolidation of Memories of Emotionally Arousing Experiences. Annu Rev Neurosci 27:1–28. doi:10.1146/annurev.neuro.27.070203.144157

42. McGaugh JL, McIntyre CK, Power AE. 2002. Amygdala Modulation of Memory Consolidation: Interaction with Other Brain Systems. Neurobiol Learn Mem 78:539–552. doi:10.1006/nlme.2002.4082

43. Mercier MR, Dubarry A-S, Tadel F, Avanzini P, Axmacher N, Cellier D, Vecchio MD, Hamilton LS, Hermes D, Kahana MJ, Knight RT, Llorens A, Megevand P, Melloni L, Miller KJ, Piai V, Puce A, Ramsey NF, Schwiedrzik CM, Smith SE, Stolk A, Swann NC, Vansteensel MJ, Voytek B, Wang L, Lachaux J-P, Oostenveld R. 2022. Advances in human intracranial electroencephalography research, guidelines and good practices. Neuroimage 260:119438. doi:10.1016/j.neuroimage.2022.119438

44. Minxha J, Adolphs R, Fusi S, Mamelak AN, Rutishauser U. 2020. Flexible recruitment of memory-based choice representations by the human medial frontal cortex. Science 368:eaba3313. doi:10.1126/science.aba3313

45. Mohan UR, Watrous AJ, Miller JF, Lega BC, Sperling MR, Worrell GA, Gross RE, Zaghloul KA, Jobst BC, Davis KA, Sheth SA, Stein JM, Das SR, Gorniak R, Wanda PA, Rizzuto DS, Kahana MJ, Jacobs J. 2020. The effects of direct brain stimulation in humans depend on frequency, amplitude, and white-matter proximity. Brain Stimul 13:1183–1195. doi:10.1016/j.brs.2020.05.009

46. Paulk AC, Zelmann R, Crocker B, Widge AS, Dougherty DD, Eskandar EN, Weisholtz DS, Richardson RM, Cosgrove GR, Williams ZM, Cash SS. 2022. Local and distant cortical responses to single pulse intracranial stimulation in the human brain are differentially modulated by specific stimulation parameters. Brain Stimul 15:491–508. doi:10.1016/j.brs.2022.02.017

47. Pedregosa F, Varoquaux G, Gramfort A, Michel V, Thirion B, Grisel O, Blondel M, Müller A, Nothman J, Louppe G, Prettenhofer P, Weiss R, Dubourg V, Vanderplas J, Passos A, Cournapeau D, Brucher M, Perrot M, Duchesnay É. 2012. Scikit-learn: Machine Learning in Python. Arxiv.

48. Peyrache A, Dehghani N, Eskandar EN, Madsen JR, Anderson WS, Donoghue JA, Hochberg LR, Halgren E, Cash SS, Destexhe A. 2012. Spatiotemporal dynamics of neocortical excitation and inhibition during human sleep. Proc Natl Acad Sci 109:1731–1736. doi:10.1073/pnas.1109895109

49. Phelps EA. 2006. Emotion and Cognition: Insights from Studies of the Human Amygdala. Annu Rev Psychol 57:27–53. doi:10.1146/annurev.psych.56.091103.070234

50. Phelps EA. 2004. Human emotion and memory: interactions of the amygdala and hippocampal complex. Curr Opin Neurobiol 14:198–202. doi:10.1016/j.conb.2004.03.015

51. Phelps EA, LeDoux JE. 2005. Contributions of the Amygdala to Emotion Processing: From Animal Models to Human Behavior. Neuron 48:175–187. doi:10.1016/j.neuron.2005.09.025

52. Qasim SE, Mohan UR, Stein JM, Jacobs J. 2023. Neuronal activity in the human amygdala and hippocampus enhances emotional memory encoding. Nat Hum Behav 7:754–764. doi:10.1038/s41562-022-01502-8

53. Quyen MLV, Bragin A, Staba R, Crépon B, Wilson CL, Engel J. 2008. Cell Type-Specific Firing during Ripple Oscillations in the Hippocampal Formation of Humans. J Neurosci 28:6104–6110. doi:10.1523/jneurosci.0437-08.2008

54. Richardson MP, Strange BA, Dolan RJ. 2004. Encoding of emotional memories depends on amygdala and hippocampus and their interactions. Nat Neurosci 7:278–285. doi:10.1038/nn1190

55. Roesler R, Parent MB, LaLumiere RT, McIntyre CK. 2021. Amygdala-hippocampal interactions in synaptic plasticity and memory formation. Neurobiol Learn Mem 184:107490. doi:10.1016/j.nlm.2021.107490

56. Romero MC, Merken L, Janssen P, Davare M. 2022. Neural effects of continuous theta-burst stimulation in macaque parietal neurons. eLife 11:e65536. doi:10.7554/elife.65536

57. Roozendaal B, McEwen BS, Chattarji S. 2009. Stress, memory and the amygdala. Nat Rev Neurosci 10:423–433. doi:10.1038/nrn2651

58. Roy AK, Shehzad Z, Margulies DS, Kelly AMC, Uddin LQ, Gotimer K, Biswal BB, Castellanos FX, Milham MP. 2009. Functional connectivity of the human amygdala using resting state fMRI. NeuroImage 45:614–626. doi:10.1016/j.neuroimage.2008.11.030

59. Rutishauser U, Ross IB, Mamelak AN, Schuman EM. 2010. Human memory strength is predicted by theta-frequency phase-locking of single neurons. Nature 464:903–907. doi:10.1038/nature08860

60. Schonhaut DR, Rao AM, Ramayya AG, Solomon EA, Herweg NA, Fried I, Kahana MJ. 2024. MTL neurons phase-lock to human hippocampal theta. eLife 13:e85753. doi:10.7554/elife.85753

61. Schwabe L, Tegenthoff M, Höffken O, Wolf OT. 2013. Mineralocorticoid Receptor Blockade Prevents Stress-Induced Modulation of Multiple Memory Systems in the Human Brain. Biol Psychiatry 74:801–808. doi:10.1016/j.biopsych.2013.06.001

62. Seabold S, Perktold J. 2010. Statsmodels: Econometric and Statistical Modeling with Python. Proc 9th Python Sci Conf 92–96. doi:10.25080/majora-92bf1922-011

63. Shoham S, Fellows MR, Normann RA. 2003. Robust, automatic spike sorting using mixtures of multivariate t-distributions. J Neurosci Methods 127:111–122. doi:10.1016/s0165-0270(03)00120-1

64. Siegle JH, Wilson MA. 2014. Enhancement of encoding and retrieval functions through theta phase-specific manipulation of hippocampus. Elife 3:e03061. doi:10.7554/elife.03061

65. Solomon EA, Kragel JE, Gross R, Lega B, Sperling MR, Worrell G, Sheth SA, Zaghloul KA, Jobst BC, Stein JM, Das S, Gorniak R, Inman CS, Seger S, Rizzuto DS, Kahana MJ. 2018. Medial temporal lobe functional connectivity predicts stimulation-induced theta power. Nat Commun 9:4437. doi:10.1038/s41467-018-06876-w

66. Solomon EA, Sperling MR, Sharan AD, Wanda PA, Levy DF, Lyalenko A, Pedisich I, Rizzuto DS, Kahana MJ. 2021. Theta-burst stimulation entrains frequency-specific oscillatory responses. Brain Stimul 14:1271–1284. doi:10.1016/j.brs.2021.08.014

67. Stein JL, Wiedholz LM, Bassett DS, Weinberger DR, Zink CF, Mattay VS, Meyer-Lindenberg A. 2007. A validated network of effective amygdala connectivity. NeuroImage 36:736–745. doi:10.1016/j.neuroimage.2007.03.022

68. Stiso J, Khambhati AN, Menara T, Kahn AE, Stein JM, Das SR, Gorniak R, Tracy J, Litt B, Davis KA, Pasqualetti F, Lucas TH, Bassett DS. 2019. White Matter Network Architecture Guides Direct Electrical Stimulation through Optimal State Transitions. Cell Reports 28:2554–2566.e7. doi:10.1016/j.celrep.2019.08.008

69. Suppa A, Huang Y-Z, Funke K, Ridding MC, Cheeran B, Lazzaro VD, Ziemann U, Rothwell JC. 2016. Ten Years of Theta Burst Stimulation in Humans: Established Knowledge, Unknowns and Prospects. Brain Stimul 9:323–335. doi:10.1016/j.brs.2016.01.006

70. Teleńczuk M, Teleńczuk B, Destexhe A. 2020. Modelling unitary fields and the single-neuron contribution to local field potentials in the hippocampus. J Physiol 598:3957– 3972. doi:10.1113/jp279452

71. Titiz AS, Hill MRH, Mankin EA, Aghajan ZM, Eliashiv D, Tchemodanov N, Maoz U, Stern J, Tran ME, Schuette P, Behnke E, Suthana NA, Fried I. 2017. Theta-burst microstimulation in the human entorhinal area improves memory specificity. Elife 6:e29515. doi:10.7554/elife.29515

72. Van der Plas M, Roux F, Chelvarajah R, Sawlani V, Staresina B, Wimber M, Rollings DT, Hanslmayr S. 2024. Characterizing neuronal and population responses to electrical stimulation in the human hippocampo-cortical network. bioRxiv 2024.11.28.625915. doi:10.1101/2024.11.28.625915

73. Vinck M, Oostenveld R, Wingerden M van, Battaglia F, Pennartz CMA. 2011. An improved index of phase-synchronization for electrophysiological data in the presence of volume-conduction, noise and sample-size bias. Neuroimage 55:1548– 1565. doi:10.1016/j.neuroimage.2011.01.055

74. Virtanen P, Gommers R, Oliphant TE, Haberland M, Reddy T, Cournapeau D, Burovski E, Peterson P, Weckesser W, Bright J, Walt SJ van der, Brett M, Wilson J, Millman KJ, Mayorov N, Nelson ARJ, Jones E, Kern R, Larson E, Carey CJ, Polat İ, Feng Y, Moore EW, VanderPlas J, Laxalde D, Perktold J, Cimrman R, Henriksen I, Quintero EA, Harris CR, Archibald AM, Ribeiro AH, Pedregosa F, Mulbregt P van, Contributors S 1 0 Vijaykumar A, Bardelli AP, Rothberg A, Hilboll A, Kloeckner A, Scopatz A, Lee A, Rokem A, Woods CN, Fulton C, Masson C, Häggström C, Fitzgerald C, Nicholson DA, Hagen DR, Pasechnik DV, Olivetti E, Martin E, Wieser E, Silva F, Lenders F, Wilhelm F, Young G, Price GA, Ingold G-L, Allen GE, Lee GR, Audren H, Probst I, Dietrich JP, Silterra J, Webber JT, Slavič J, Nothman J, Buchner J, Kulick J, Schönberger JL, Cardoso JV de M, Reimer J, Harrington J, Rodríguez JLC, Nunez-Iglesias J, Kuczynski J, Tritz K, Thoma M, Newville M, Kümmerer M, Bolingbroke M, Tartre M, Pak M, Smith NJ, Nowaczyk N, Shebanov N, Pavlyk O, Brodtkorb PA, Lee P, McGibbon RT, Feldbauer R, Lewis S, Tygier S, Sievert S, Vigna S, Peterson S, More S, Pudlik T, Oshima T, Pingel TJ, Robitaille TP, Spura T, Jones TR, Cera T, Leslie T, Zito T, Krauss T, Upadhyay U, Halchenko YO, Vázquez-Baeza Y. 2020. SciPy 1.0: fundamental algorithms for scientific computing in Python. Nat Methods 17:261–272. doi:10.1038/s41592-019-0686-2

75. Yoo HB, Umbach G, Lega B. 2021. Neurons in the human medial temporal lobe track multiple temporal contexts during episodic memory processing. NeuroImage 245:118689. doi:10.1016/j.neuroimage.2021.118689

76. Youssef D, Wittig JH, Jackson S, Inati SK, Zaghloul KA. 2023. Neuronal Spiking Responses to Direct Electrical Microstimulation in the Human Cortex. J Neurosci 43:4448–4460. doi:10.1523/jneurosci.1666-22.2023

77. Zheng J, Anderson KL, Leal SL, Shestyuk A, Gulsen G, Mnatsakanyan L, Vadera S, Hsu FPK, Yassa MA, Knight RT, Lin JJ. 2017. Amygdala-hippocampal dynamics during salient information processing. Nat Commun 8:14413. doi:10.1038/ncomms14413

78. Zheng J, Stevenson RF, Mander BA, Mnatsakanyan L, Hsu FPK, Vadera S, Knight RT, Yassa MA, Lin JJ. 2019. Multiplexing of Theta and Alpha Rhythms in the Amygdala-Hippocampal Circuit Supports Pattern Separation of Emotional Information. Neuron 102:887–898.e5. doi:10.1016/j.neuron.2019.03.025

