## Supplementary figures and images for "Human single-neuron activity is modulated by intracranial theta burst stimulation of the basolateral amygdala"

### Figure S1

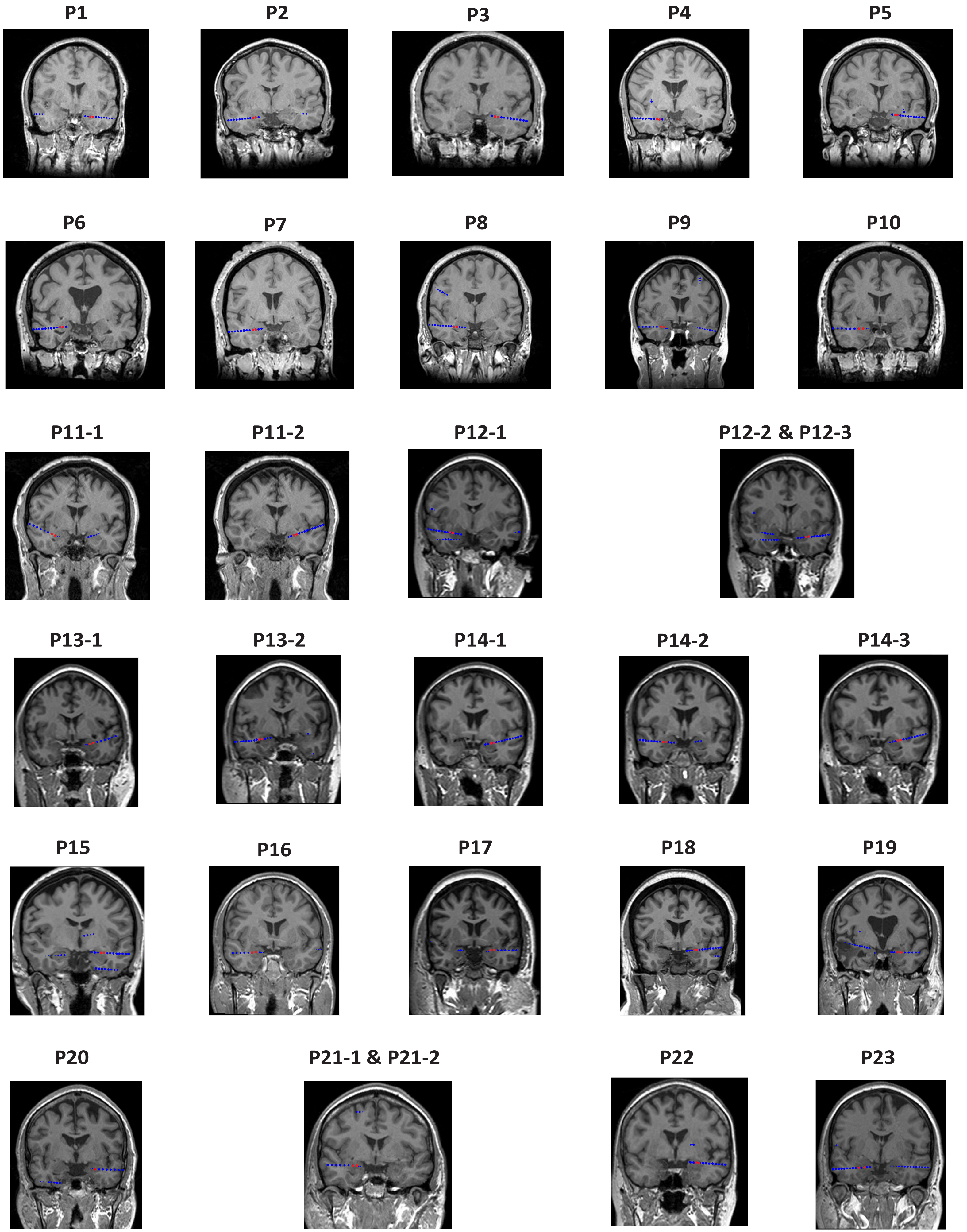

### Figure S2

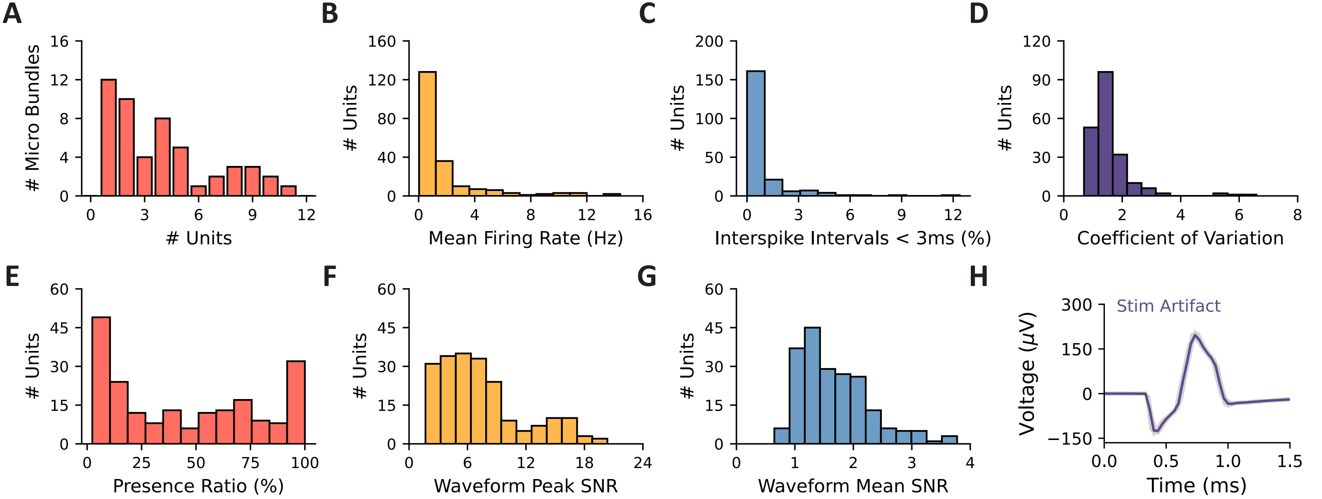

### Figure S3

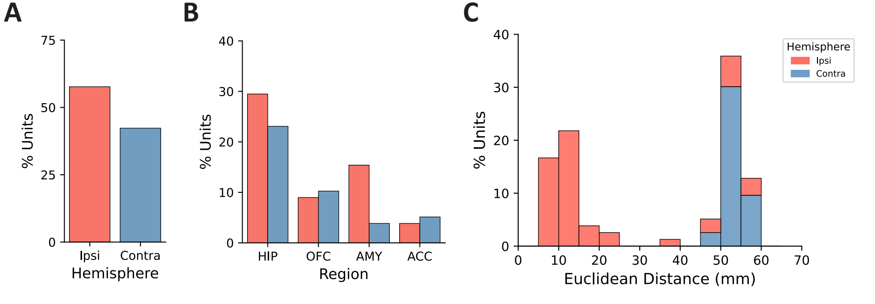

### Figure S4

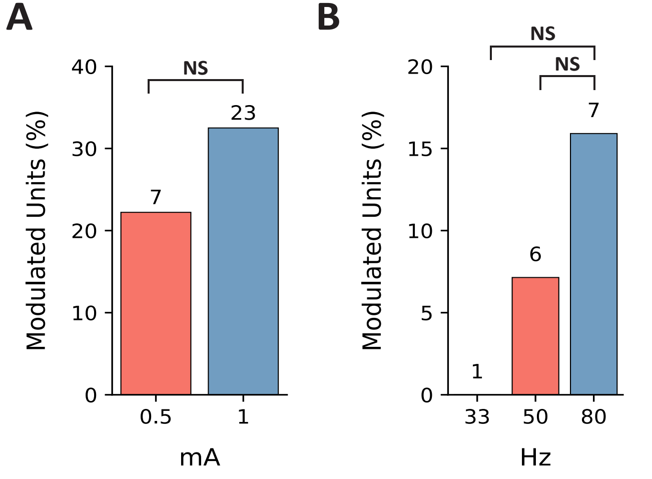

### Figure S5

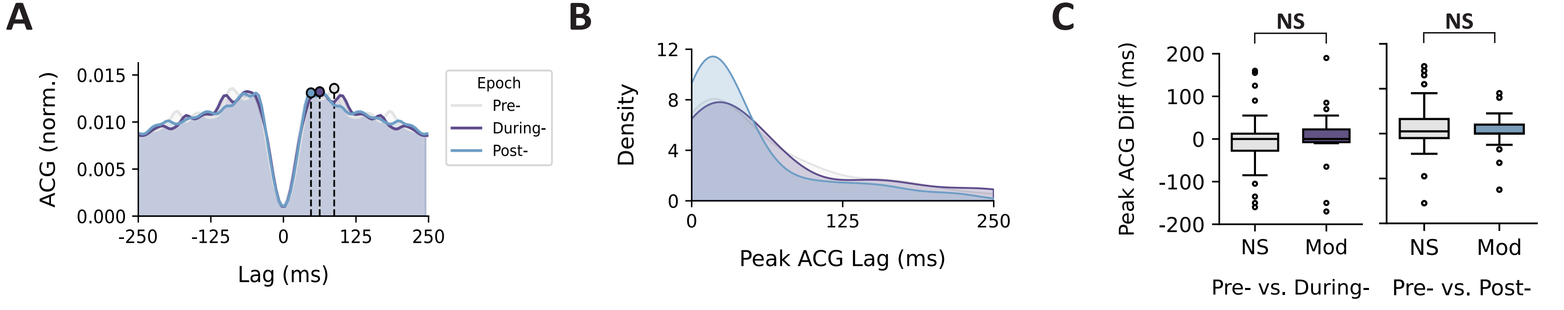

### Figure S6

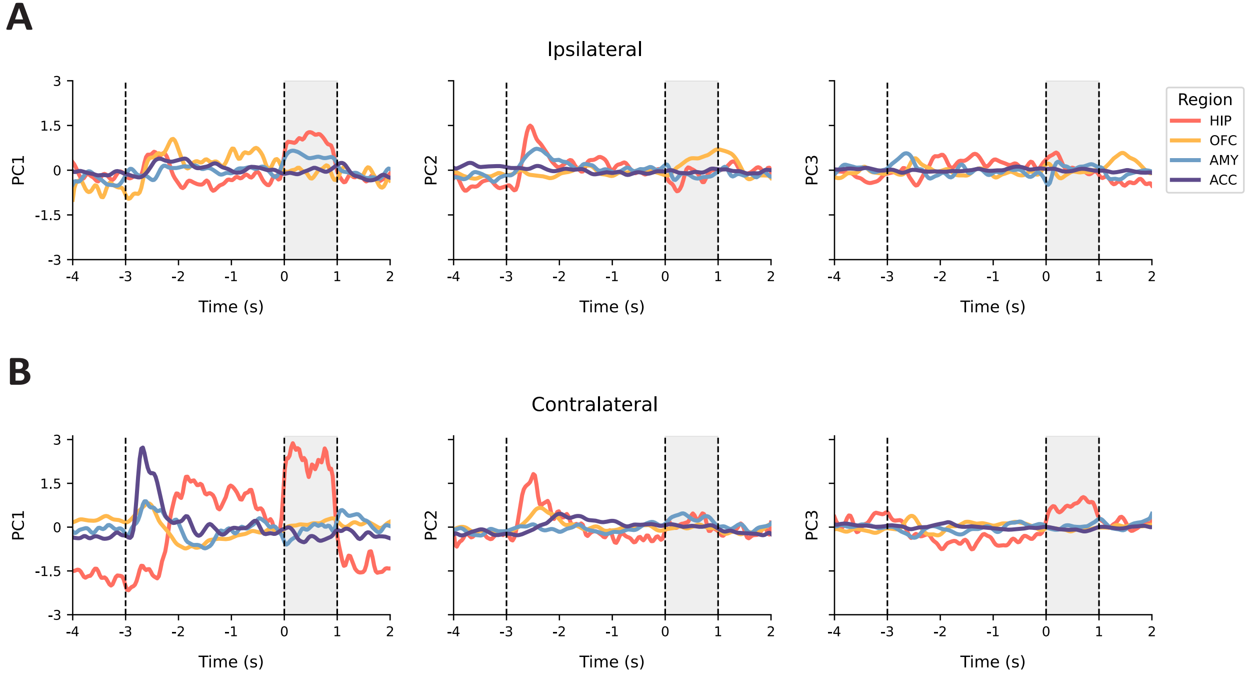

### Figure S7

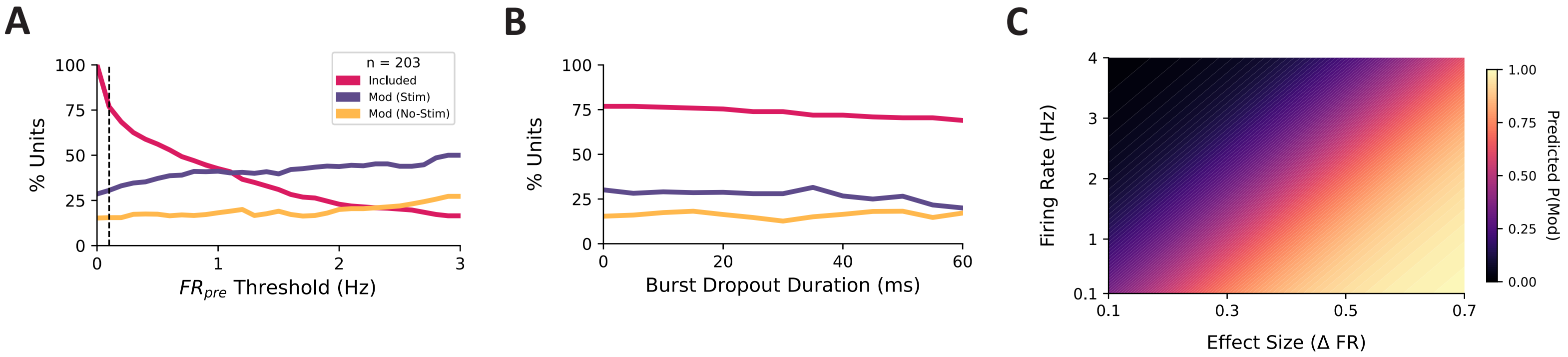

### Figure S8

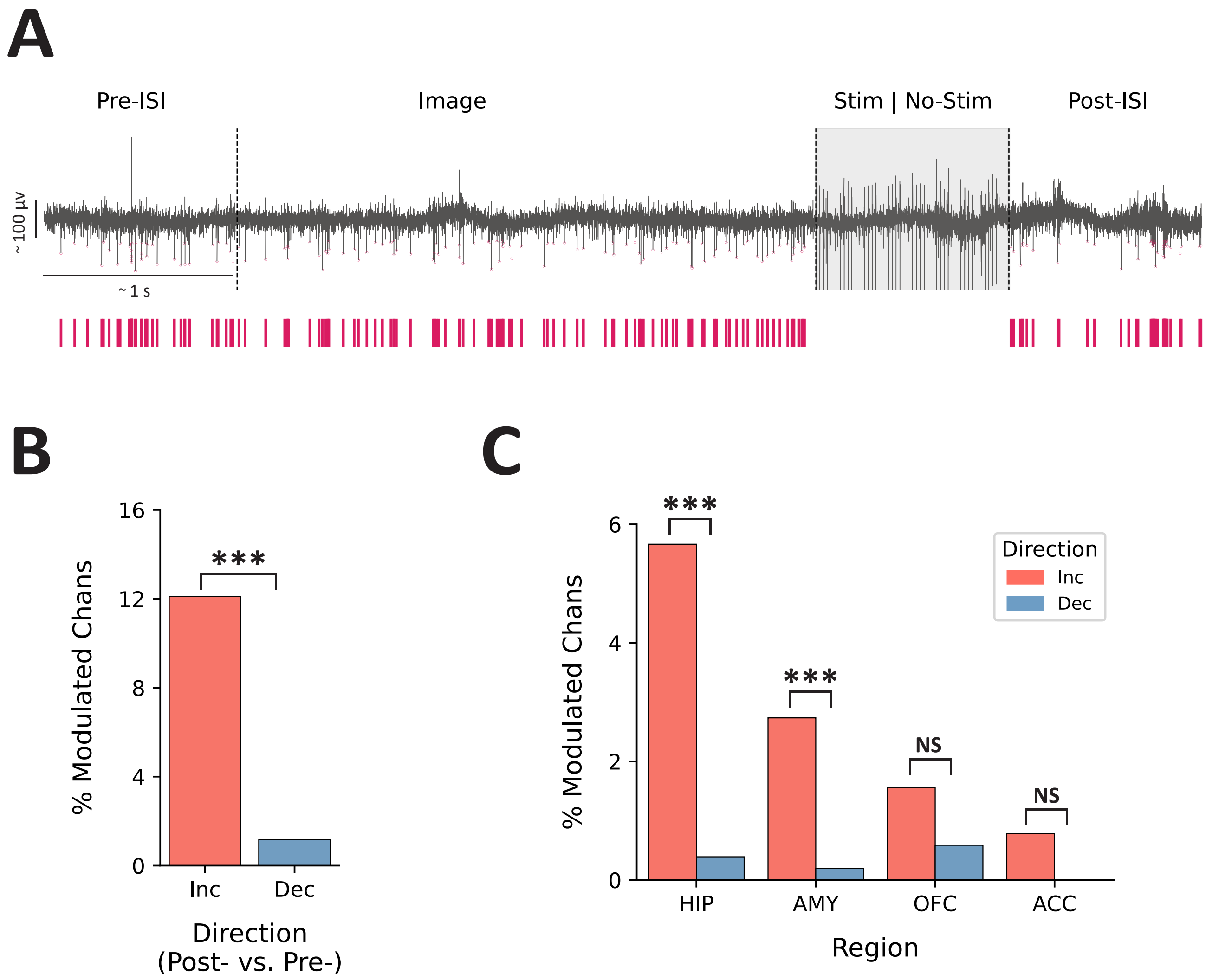

### Figure S9

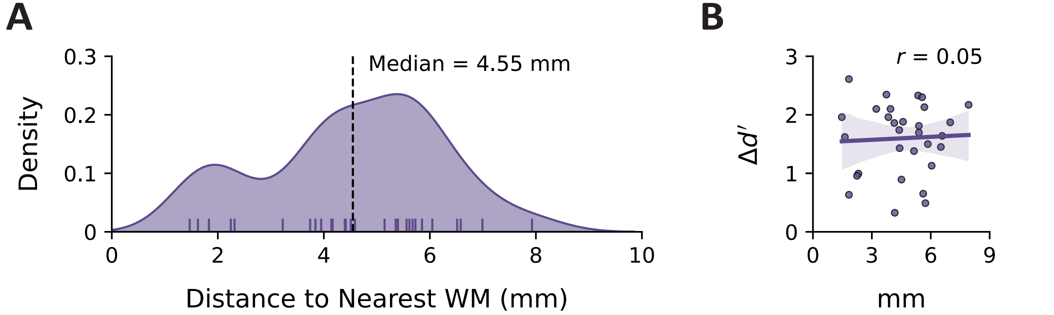
