## Supplementary material for "Human single-neuron activity is modulated by intracranial theta burst stimulation of the basolateral amygdala": Table S1

**Supplemental Table 1. Patient demographics and clinical characteristics.**

| **Patient** | **Age**  **(yrs)** | **Sex** | **Race** | **# Sessions** | **# Leads (micros)** | **# Units (L, R)** | **Seizure Onset Zone** |
| --- | --- | --- | --- | --- | --- | --- | --- |
| P1 | 40 | F | White | 1 | 9 (3) | 14 (6, 8) | BL hippocampi |
| P2 | 35 | F | White | 1 | 8 (2) | 7 (0, 7) | R hippocampus and amygdala |
| P3 | 44 | F | White | 1 | 10 (3) | 7 (7, 0) | L hippocampus and temporooccipital junction |
| P4 | 19 | F | White | 1 | 8 (3) | 12 (0, 12) | R medial temporal lobe |
| P5 | 23 | F | American Indian or Alaska Native | 1 | 6 (3) | 3 (1, 2) | L medial temporal lobe |
| P6 | 66 | F | White | 1 | 10 (3) | 5 (5, 0) | BL hippocampi and amygdalae |
| P7 | 44 | F | White | 1 | 8 (3) | 21 (12, 9) | L hippocampus |
| P8 | 28 | F | White | 1 | 8 (3) | 2 (0, 2) | R anterior temporal lobe |
| P9 | 42 | F | White | 1 | 13 (3) | 20 (13, 7) | L hippocampus and amygdala |
| P10 | 40 | F | White | 1 | 9 (3) | 5 (0, 5) | L medial temporal lobe |
| P11 | 38 | F | White | 2 | 18 (2) | 14 (0, 14) | BL hippocampi, R temporal pole, L temporal gyrus |
| P12 | 45 | F | White | 3 | 12 (2) | 17 (8, 9) | BL hippocampi |
| P13 | 40 | M | White | 2 | 11 (2) | 8 (8, 0) | L hippocampus and entorhinal cortex |
| P14 | 37 | M | White | 3 | 14 (2) | 16 (16, 0) | R hippocampus and amygdala |
| P15 | 34 | F | White | 1 | 13 (2) | 6 (6, 0) | L hippocampus and amygdala |
| P16 | 37 | M | White | 1 | 18 (2) | 6 (0, 6) | BL hippocampi |
| P17 | 24 | F | Black or African American | 1 | 10 (2) | 11 (11, 0) | R hippocampus and entorhinal cortex |
| P18 | 41 | F | Black or African American | 1 | 15 (2) | 5 (0, 5) | BL anterior temporal pole |
| P19 | 38 | M | White | 1 | 13 (2) | 11 (11, 0) | R hippocampus, orbitofrontal cortex, and middle temporal gyrus |
| P20 | 41 | M | White | 1 | 11 (2) | 1 (1, 0) | R temporal lobe |
| P21 | 22 | M | White | 2 | 17 (2) | 5 (0, 5) | R frontotemporal region |
| P22 | 60 | F | White | 1 | 17 (1) | 2 (2, 0) | L middle frontal gyrus and superior frontal gyrus |
| P23 | 40 | F | White | 1 | 13 (2) | 5 (0, 5) | L hippocampus and amygdala |

L = left, R = right, BL = bilateral.
